# PhAT: A flexible open-source GUI-driven toolkit for photometry analysis

**DOI:** 10.1101/2023.03.14.532489

**Authors:** Kathleen Z. Murphy, Eyobel Haile, Anna McTigue, Anne F. Pierce, Zoe R. Donaldson

## Abstract

Photometry approaches detect sensor-mediated changes in fluorescence as a proxy for rapid molecular changes within the brain. As a flexible technique with a relatively low cost to implement, photometry is rapidly being incorporated into neuroscience laboratories. While multiple data acquisition systems for photometry now exist, robust analytical pipelines for the resulting data remain limited. Here we present the <u>Ph</u>otometry <u>A</u>nalysis <u>T</u>oolkit (PhAT) - a free open source analysis pipeline that provides options for signal normalization, incorporation of multiple data streams to align photometry data with behavior and other events, calculation of event-related changes in fluorescence, and comparison of similarity across fluorescent traces. A graphical user interface (GUI) enables use of this software without prior coding knowledge. In addition to providing foundational analytical tools, PhAT is designed to readily incorporate community-driven development of new modules for more bespoke analyses, and data can be easily exported to enable subsequent statistical testing and/or code-based analyses. In addition, we provide recommendations regarding technical aspects of photometry experiments including sensor selection and validation, reference signal considerations, and best practices for experimental design and data collection. We hope that the distribution of this software and protocol will lower the barrier to entry for new photometry users and improve the quality of collected data, increasing transparency and reproducibility in photometry analyses.

Basic Protocol 1: Software Environment Installation
Basic Protocol 2: GUI-driven Fiber Photometry Analysis
Basic Protocol 3: Adding Modules

## INTRODUCTION

Fiber photometry is a method for recording bulk fluorescence changes in the brain at subsecond timescales, often employed in behaving animals. Fiber photometry has gained substantial popularity in neuroscience labs since original reports detailing the technique were published in 2014 (Gunaydin et al., 2014). Several factors have contributed to this popularity, including an expanding toolbox of sub-second resolution fluorescent biosensors that detect a range of substances within the brain (O’Banion and Yasuda, 2020; Akerboom et al., 2012; Marvin et al., 2013; Sun et al., 2018; Feng et al., 2019), the relative ease of implementation among labs that already perform intracranial surgery, and its relatively low cost. In addition, because sensor delivery can be achieved via viral vector infusions and small diameter ferrules, photometry can be relatively easily implemented in less commonly employed laboratory species where transgenic technologies and/or more invasive approaches may be highly challenging.

Despite the widespread adoption of fiber photometry, the subsequent analysis remains challenging for many labs. Here we introduce the graphical user interface (GUI)-based **<u>Ph</u>**otometry **<u>A</u>**nalysis **<u>T</u>**oolkit (**PhAT**), which enables rapid examination and analysis of fiber photometry data in relation to behavior or other metrics. This modular, python-based toolkit enables tremendous flexibility for users to analyze data within the GUI, which requires no coding skills. PhAT adds a few key features to an existing set of fiber photometry analysis softwares, such as GuPPY (Sherathiya et al., 2021) and pMAT (Bruno et al., 2021). Its modular object-oriented design enables straightforward addition of new modules, making this software a solid foundation for the python community to create and publish new analyses and functionality. It also includes multiple approaches for signal normalization and motion correction that can be evaluated and chosen based on the relevant attributes within the collected data. Finally, it can flexibly incorporate data from multiple time-stamped data streams and includes an import option for working with standard Neurophotometrics and BORIS data outputs.

Below we outline some considerations for conducting fiber photometry experiments that will help optimize data quality and ease of analysis. We then outline protocols for installing (new users) and updating (current users) the PhAT software and python environment. The following protocol details how to interact with the GUI and use each of the current modules to analyze and evaluate data. The last protocol describes the process for adding new functionality to the software either through the GUI or using the jupyter notebook.

## STRATEGIC PLANNING

As with any experiment, a successful outcome depends on careful consideration of experimental design that incorporates the strengths and limitations of the technologies being employed. The below considerations are not meant to comprehensively address all technical aspects of working with biosensors in fiber photometry applications. Rather this is intended to serve as a starting point for successful implementation with reference to additional available resources indicated below.

### Choosing a sensor

There now exist several fluorescent molecular sensors designed to measure Ca^2+^ activity as a metric of neuronal activity or measure extracellular levels of various signaling molecules (dopamine, serotonin, oxytocin, vasopressin, glutamate, GABA, etc.) Such sensors often employ a circularly permuted GFP linked to either a G-protein coupled receptor (GPCR; as in dLight and GRAB-type sensors) (Patriarchi et al., 2018; Sun et al., 2018; Feng et al., 2019; Wan et al., 2020) or a binding protein (as in GCaMP and -snfr’s) (Akerboom et al., 2012; Marvin et al., 2013). Less commonly employed are FRET-based fluorescent sensors (Jones-Tabah et al., 2022). Each of these has advantages and disadvantages, but the specific sensor employed may ultimately depend on practical considerations related to availability and localized expertise. The majority of these sensors are designed for interrogation via green fluorescence, but a handful now exist that use red-shifted excitation, allowing for detection of two spectrally-distinct sensors within a given brain region (Akerboom et al., 2013; Patriarchi et al., 2020).

### Identifying a reference signal

Reference signals provide a means to detect and potentially subtract out motion artifacts. For systems where more than one wavelength can be collected, the choice of sensor will guide subsequent choice of a reference signal. For well-established sensors, there is often a known isosbestic point at which the fluorescence emission of the sensor is signal independent. For GCaMP6m, the isosbestic point is 410 nm, and thus many systems are built to assess 405 - 415 nm as the reference signal (Feshki et al., 2020; Martianova et al., 2019; Chen et al., 2013). However, for other sensors, 405-415 nm may not represent the isosbestic point, and collection of data at that wavelength serves as a poor reference signal. For example, if you excite GRAB_DA_ (isosbestic ~ 440 nm) at 415 nm, it will be less bright when in the DA bound state than in the unbound state, creating an inversion of the 470nm signal (Sun et al., 2020). If 415 nm fluorescence is used as a reference to remove motion artifacts, it will instead non-linearly amplify the 470 nm signal and less effectively reduce motion artifacts, impairing the interpretability of the data. In instances where the sensor’s isosbestic point is not well delineated or the system does not allow for recording at the appropriate wavelength, the most conservative path is to use a second, spectrally distinct and signal-independent fluorophore, such as mCherry (Pierce et al., 2022). In a subset of fiber photometry systems (such as Amuza), no reference signal is queried, and in those instances, it is essential to include a control group of animals expressing only a corresponding signal-independent fluorophore (e.g. YFP/GFP, mCherry/tdTomato, or an inactive mutant sensor) to ensure that observed fluorescent changes are not due to motion (Matias et al., 2017; Gunaydin et al., 2014; Wan et al., 2020).

### Experimental Design Considerations

Fiber photometry can be used to measure relative changes in fluorescence within an animal during a recording session. It cannot be interpreted as an absolute measure of a molecule’s activity in a region, and therefore raw values should never be compared between animals. Inter-animal variation can result from differences in sensor expression and ferrule placement relative to sensor-expressing cells. Of note, fluorescence intensity and signal to noise ratio can also vary within animals due to several factors. To decrease day-to-day variation in recordings, we recommend the following: 1) Confirm the time course over which your sensor expression plateaus in your brain region of interest and commence recordings once expression has reached a steady state. For this, the promoter driving the sensor can be an important factor. 2) Keep light power consistent across recording days. 3) Pay attention to fiber-optic connectivity; gaps between the patch cable and the ferrule will result in changes to the detected signal.

As such, within-animal and ideally within-trial designs are best for examining event-related changes in signal intensity. When comparing measures between recording sessions or between animals, it is imperative to use relative measures such as percent changes or changes in z score to account for the variability described above (Li et al., 2019).

As outlined below, motion correction approaches are not foolproof; we recommend running controls that express a signal-insensitive fluorophore (Matias et al., 2017; Gunaydin et al., 2014). In some cases, there exist control versions of sensors developed for this purpose (Wan et al., 2020; Feng et al., 2019).

### Reducing motion artifacts

The ability to record neural activity from active animals is one of the strengths of fiber photometry. However, movement can introduce artifacts to your signal. While motion artifacts can be corrected for post hoc by using a reference channel (Lerner et al., 2015; Akerboom et al., 2012; Girven and Sparta, 2017), such motion-correction strategies have limitations. Taking steps to reduce motion artifacts before and during data collection is important for optimizing the quality of your data.

Motion artifacts originate from two sources. First, bending of the photometry tether and/or tension on the tether can contribute to motion artifacts. These can be reduced by choosing recording arenas that reduce the chances of bending and tugging and supporting the weight of the fiber by hanging it from a higher location or a helium balloon. In addition, using a commutator can help alleviate stress on the fiber optic cable, but this can decrease signal. Thus, a commutator is not advisable for applications in which low signal is expected, such as certain sensors or when recording CA^2+^ activity in neuronal terminals. Second, motion artifacts can occur when the implanted ferrule shifts relative to the brain. Making the fiber as stable as possible will help reduce these motion artifacts and decrease chances of the fiber completely dislodging before the end of the experiment. Ways to ensure stability include making sure the skull is dry and clean of blood and tissue prior to adhesive application, scoring the skull lightly with a scalpel or chemical etchant, using a stronger cement or adhesive, and maintaining excellent aseptic technique and using peri/postoperative antibiotics and anti-inflammatory drugs to reduce infection risk and inflammation. Finally, motion artifacts are often more pronounced in deeper brain regions where the end of the ferrule is farther from the skull, and these can be ameliorated by adding 1 - 2 wires affixed to the sides of the ferrule that extend beyond its end and help anchor the tissue around the base of the ferrule.

### Optimizing Fluorescence Collection

Most fiber photometry systems allow for control of the excitation light source power. Increasing the power will increase your signal to noise and may be useful or necessary when working with low signal to noise sensors, recording from cell projection terminals, or in regions with low signal. However, increasing the power of your excitation light source will also increase photobleaching and may even cause tissue damage or cell death, especially when recording at high frame rates over long periods of time (Akerboom et al., 2012; Girven and Sparta, 2017).

In fiber photometry, the time resolution of your data is limited by the dynamics of your sensor and the frame rate of your acquisition system. Setting your frame rate to be twice as fast as your sensor dynamics will give you the highest possible time resolution. For example, GRAB_DA_ has a rise time of 0.08 sec, if you take a frame every 0.04 sec (25Hz), you will be able to detect all real rises in your sensor; increasing your frame rate will not increase your time resolution. However, depending on the design of your experiment and the temporal dynamics you wish to capture, your data may not require the highest temporal resolution. In such instances, decreasing the frame rate can help combat photobleaching and tissue damage due to high light powers.

When recording at multiple wavelengths, each light source can be turned on sequentially or they can all be turned on simultaneously. While the sequential option will reduce the highest possible frame rate, we always recommend this option because simultaneous excitation at multiple wavelengths greatly increases the chance of signal bleed-through.

### Synchronizing data streams

Fiber photometry is often collected alongside other data, such as behavioral video recordings or devices that detect specific actions, such as lickometer strokes, nose-pokes, or lever-pressing. To accurately align neural data with data from other sources, it is important to be certain that your data streams are aligned properly.

The easiest way to align datastreams is to use a data acquisition software such as BONSAI, to collect all your data streams using a shared clock. When this is not possible you can align your data streams post hoc. For example, if all instruments are alignedto a universal clock, then the timestamps can be aligned. Alternately, a flashing light, that is time stamped with the same clock as your fiber photometry data, can be added to your behavior video to serve as a synchronization cue. It is very important when collecting fiber photometry data, videos, or other sequential data that each frame has a timestamp, because even if your frame rate is very regular, dropped frames are common and can cause large shifts in time alignment throughout a session if you extrapolate from the time of the first frame.

Our software allows for two format options when importing behavior data. The BORIS format assumes that the zero time in your behavior data corresponds to the first value in your fiber photometry data file. The alternative format assumes that the first timestamp corresponds with the first value in your fiber photometry data file.

### Validating your sensor

While it is necessary to validate each sensor in each region you plan to employ it in, we recommend initially testing a new-to-your-lab sensor in a region that is easy to surgically target and/or has documented robust dynamics for the molecule you plan to detect (e.g. we validated our GRAB_DA_ dopamine sensor in the nucleus accumbens). Resendez et al provide recommendations to optimize viral expression of your sensor (Resendez et al., 2016). Briefly, considerations include: optimizing titer and injection volume to ensure expression in your region of interest without ectopic expression or cell death. Of note, viral expression beyond your region of interest is not necessarily a major concern as fluorescence changes will be detected only within the region proximal to the end of the implanted ferrule.

Identifying optimal stereotactic coordinates for your brain region of interest may require some trial and error. The most expeditious way we have found to assess stereotactic coordinates is to implant a ferrule and immediately perfuse the animal to assess location. For viral spread, we generally recommend waiting 3-4 weeks for most vectors if assessing somatic expression and 6+ weeks for expression at terminals. For sensors with poorer signal-to-noise dynamics, consider using fluorescence-guided ferrule implantation to ensure ferrule placement within the bulk of your fluorescence. With this approach, you will inject your viral vector and then wait 2 - 3 weeks for expression before lowering the ferrule into place while simultaneously recording fluorescence values with your fiber photometry system, affixing the ferrule when it reaches the intended coordinates, and you observe a detectable increase in fluorescence.

Once you have optimized surgical procedures, you will need to validate your sensor. In addition to determining that you can detect fluorescence increases and decreases independent of motion artifacts (see support protocol 2a), we also recommend an additional step to block, increase, and/or decrease the molecule that the sensor is designed to detect and examine subsequent changes in fluorescence. This is particularly important when working with less commonly employed sensors. The following are three strategies for assessing sensor activity in vivo:

i. Pharmacological blockade of sensor function: the activity of sensors designed to detect neuromodulators/hormones, and fluorescence changes can be effectively blocked through addition of a molecule that prevents binding of the target molecule to the sensor. For instance, fluorescence changes from GRAB_DA_ are blocked by the dopamine D2 receptor antagonist, eticlopride (Sun et al., 2020). These are best employed in an intra-animal design that compares fluorescence in vehicle versus drug conditions, ideally including a behavior/event that is known to elicit release of your neuromodulator of interest.
ii. Pharmacological manipulation of your target system: Alternatively, you can manipulate release or neural activity and assess subsequent changes in fluorescence. One straightforward, if indirect method for decreasing fluorescence is to record from deeply anesthetized animals in which most neural function is quiet. Conversely, pharmacological approaches can be used to elicit neural activity (for instance via seizure induction) or stimulate release and/or synaptic accumulation of your neuromodulator of interest (for instance cocaine for dopamine, MDMA for serotonin) (Feng et al., 2019; Patriarchi et al., 2020). As this approach is likely to lead to changes on longer timescales, it is important to consider the effects of sensor photobleaching. These systemic manipulations often increase or decrease motion in the same direction as neural activity; therefore, it is important to detect changes in the mean fluorescence of your signal overtime as opposed to increases or decreases in the fluctuations of the fluorescence.
iii. Optogenetic activation/inactivation: Similarly, you can assess whether optogenetic manipulation of your target system results in a corresponding change in fluorescence from your sensor. For instance, optogenetic VTA activation or inhibition should increase or decrease GRAB_DA_ fluorescence, respectively, in the nucleus accumbens. A word of caution: Many sensors are excited using a wavelength that activates many optogenetic actuators, so if you decide to optogenetically manipulate terminals in the same brain region as your sensor, you should use a spectrally (typically red) shifted opsin (Akerboom et al., 2013; Feng et al., 2019; Patriarchi et al., 2020).

## Basic Protocol 1: Software and Environment Installation

This protocol includes all steps necessary to install the software and any necessary dependencies. It provides parallel instructions for Mac, Linux and Windows users. Our installation method utilizes a virtual environment to ensure that there are no conflicting issues with any existing Python dependencies. We have instructions to install the GUI using either Anaconda or PIP/PyPI. We recommend using Anaconda to utilize the Jupyter Notebook for ease of use and increased flexibility (inline error handling, modularity, further analysis of created objects, etc).

### Materials

1. Mac, Linux or Windows Computer System
2. Python Version 3.9 or newer installed
  - https://wiki.python.org/moin/BeginnersGuide/Download
3. Anaconda OR PIP/PyPI installed
  - If you plan to use Anaconda: https://docs.anaconda.com/anaconda/install/
  - If you plan to use PIP/PyPI: https://pip.pypa.io/en/stable/installation/
4. FiberPho GUI
  - https://github.com/donaldsonlab/PhAT

### Protocol steps

1. Download Code
  a. Navigate to https://github.com/donaldsonlab/PhAT
  b. Click on the green button labeled “Code” located at the top right corner of the repository, then click on “Download ZIP” (Ensure that this file is saved locally on your device i.e. not on any cloud environments).
  c. Locate the downloaded zip file on your device and place it somewhere convenient to easily navigate to it again. Avoid cloud storage.
  d. Unzip the file by right clicking on it and selecting unzip or use an unzipping utility (e.g. WinRAR on Windows systems).
  e. Take note of the FiberPho_Main folder location (folder path needed later).
    i. Mac/Unix: Right click on the folder, Hold the Option key, and copy “PhAT” as Pathname.
    ii. Windows: Right click on the folder, select Properties, and take note of the text written next to Location on your computer, this is the folder’s path.
2. Create Virtual Environment
  - Using Anaconda (Option 1: Recommended)
    a. Open a new terminal window (Mac/Unix) or Anaconda Prompt (not Anaconda Navigator) (Windows).
    b. Navigate to the location of the “PhAT” folder (noted from Step 1C).
      i. Type the following command, instead typing your folder path within the brackets: “cd [path_to_PhAT_folder]”. Then hit enter.
      ii. Ex. cd Desktop/DonaldsonLab/PhAT
    c. Create a virtual environment and give it a name (e.g. “my_gui_env”) with the following command.
      i. “conda create -n [your_env_name] python=[version] pip”. Then hit enter.
      ii. Ex: conda create -n my_gui_env python=3.9 pip
    d. Activate the virtual environment.
      i. “conda activate [your_env_name]” Then hit enter.
      ii. Ex: conda activate my_gui_env
    e. Execute the following commands to install dependencies.
      i. Type “pip list”. Then hit enter.
        ■ No dependencies should be present since this is a new environment.
      ii. Type “pip install -r requirements.txt”. Then hit enter.
      iii. Type “pip list”. Then hit enter
        ■ All necessary dependencies should now be installed.
  - Using PIP/PyPI (Option 2)
    a. Open a new terminal window (command prompt for Windows)
    b. Navigate to the location of the “PhAT” folder (noted from Step 1C).
      i. Type the following command, instead typing your folder path within the brackets: “cd [path_to_PhAT_folder]”. Then hit enter.
      ii. Ex: cd Desktop/DonaldsonLab/PhAT
    c. Create a virtual environment and give it a name (e.g. “my_gui_env”) using one of the following commands.
      i. Mac/Unix: “python3 -m venv [your_env_name]”. Then hit enter.
      ii. Windows: “py -m venv [your_env_name]”. Then hit enter.
    d. Activate the virtual environment.
      i. Mac/Unix: “source [your_env_name]/bin/activate”. Then hit enter.
      ii. Windows: “.\[your_env_name]\Scripts\activate”. Then hit enter.
    e. Execute the following commands to install dependencies.
      i. Type “pip list”. Then hit enter.
        ■ No dependencies should be present since this is a new environment.
      ii. Type “pip install -r requirements.txt”. Then hit enter.
      iii. Type “pip list”. Then hit enter.
        ■ All necessary dependencies should now be installed.

## Alternative Protocol 1: Software and Environment Update

This protocol describes how to update your software and environment for users that have already completed an initial installation.

### Materials

1. Mac, Linux or Windows Computer System
2. Previous version of PHAT installed
  - See basic protocol 1
3. FiberPho GUI
  - https://github.com/donaldsonlab/PhAT

### Protocol steps

1. Updating Software and environment (Returning Users)
  a. Repeat step 1 and replace the old version of the “PhAT” folder with the most recent version.
  b. Open a new terminal window (Mac/Unix) or Anaconda prompt (Windows).
  c. Navigate to the location of the “PhAT” folder (noted from Step 1C).
    i. Type the following command, instead typing your folder path within the brackets: “cd [path_to_PhAT_folder]”. Then hit enter.
    ii. Ex: cd Desktop/DonaldsonLab/PhAT
  d. Activate the virtual environment.
    i. Anaconda: “conda activate [your_env_name]”. Then hit enter.
    ii. PIP and Mac/Unix: “source [your_env_name]/bin/activate”. Then hit enter.
    iii. PIP and Windows: “.\[your_env_name]\Scripts\activate. Then hit enter.
  b. Execute the following commands to update dependencies.
    i. Type “pip install -r requirements.txt”. Then hit enter.

## Basic Protocol 2: GUI-driven Fiber Photometry Analysis

PhAT is designed for user-friendly flexible analysis of fiber photometry and behavioral data through a graphical user interface (GUI). Data from each fiber is imported and saved as an object to allow for visualization and analysis. This can be performed on single or multiple channels and collection sites (i.e. ferrules) simultaneously allowing for cross-region, and cross-animal analyses. The GUI contains multiple cards (see Table 1) that each have a distinct function. Using these cards, the user can normalize traces, analyze fluorescent signals relative to behavior, and examine relationships across traces. Implementation of each of these cards is optional and independent. For instance, a user can examine the relationship between two traces (e.g. the Pearson correlation coefficient) without normalizing their data or importing behavioral information. No internet connection is needed for these steps.

**Table 1:** Glossary. Here we define some terms that will be used regularly throughout our protocols.

|  |  |
| --- | --- |
| **Channel** | Denotes fluorescence intensity data collected from a specific wavelength. Most acquisition systems provide data from one to three channels/wavelengths. |
| **Trace** | A time series of fluorescence data, which can be plotted as a continuous line with time on the x-axis and fluorescence intensity on the y-axis as in Fig 4-8. Traces can be plotted from any channel in which fluorescence data was collected and can include raw, normalized, or motion corrected data. |
| **Signal** | Refers to fluorescent information collected from the excitation wavelength of your sensor of interest. For instance, for GCaMP, this would be the trace collected from the 470 nm (or similar) wavelength. The *raw signal trace* will include deflections that represent both true sensor-mediated changes in fluorescence and those introduced from motion artifacts. |
| **Reference** | Refers to fluorescent information collected at a signal independent excitation wavelength. See Identifying a reference for more information |
| **Linearization** | Both signal and reference are linearized to adjust the trace for photobleaching of the fiber optic and fluorescent sensor (Fig 4a/b). |
| **Motion-corrected** | Following normalization, the reference signal is used to remove motion artifacts from the raw signal, yielding an adjusted trace that can be most accurately interpreted in relation to behavior or other variables (Fig 4c/d). |
| **Normalization** | A flexible process that can combines linearization and motion-correction to produce a $\Delta F/F$ trace from your raw signal trace. |
| **Object** | A compilation of all the information and variables associated with one recording from one fiber, including all recording wavelengths and matched behavioral or other data. For a list of data and variables that can be included in an object see table 2. |
| **Card** | A component contained in an individual box within the GUI and used to do a specific task or analysis. |
| **Widget** | A subcomponent in a card that allows user input. These include the choose file button, drop down menus and text input boxes. |

### Materials

1. Fiber photometry data in a .csv file. The GUI accepts two options.
  - **Option 1: Neurophotmetrics (NPM) format** The first is the standard NPM output file (Fig 1a). To use this format you will need columns titled “Timestamp” and “LEDstate”. The fluorescence data will be in a series of green (G) and red (R) columns and will be interleaved based on the values in the “LEDstate” column which can be decoded using the table in the NPM FP3002 manual pg. 55 (https://neurophotometrics.com/documentation). The first G and/or R column will correspond to fiber 1, the second to fiber 2 and so on.
  - **Option 2: Alternative format** The alternative format works with non-interleaved data (Fig 1b). It must have a time column labeled “Timestamp” with data in seconds. Fluorescence data must be in any combination of columns titled: “Green”, “Red”, and “Isosbestic”. You must have at least one fluorescence data column and can have up to three. Any columns with names besides these four keywords (“Timestamp”, “Green”, “Red”, “Isosbestic”) will be ignored by the software. You will need a separate .csv for each fiberoptic within a recording session.
2. (Optional) Behavior data in a .csv file. The GUI accepts two options.
  - **Option 1: BORIS format** The BORIS format is automatically compatible with the BORIS tabular csv output (Fig 2a). To obtain this, follow these steps in the BORIS software: Observations → export events → tabular events → save as csv [*not tsv]. Although the output will work as is, the only necessary features are three columns labeled “Behavior”, “Status” and “Time” (Fig 2b). The “Behavior” column has the name of each behavior. The “Time” column has the time in seconds. And the “Status” column has the word “POINT” for discrete events (lever press, etc), or if the behavior lasts for some length of time, the word “START” and the word “STOP” for the beginning and end of a behavior bout, respectively. The order of the rows and columns does not matter but each “START” row must have a corresponding “STOP” row for that behavior. ***Important***: Time zero in your video/behavior data must correspond to the first value in your fiber photometry data file.
  - **Option 2: Alternative format** The alternative format must have a “Time” column in ms, sec, or min and columns titled for each behavior examined (Fig 2c/d). Each behavior column must consist of values assigned to indicate when a behavior occurred/did not occur, respectively (e.g. 0/1 or yes/no)(Fig 2c). While the behavior occurring value can change (e.g. 1,2,3.. or start, ongoing, end), there must only be one value indicating that a behavior is not occurring (Fig 2d). The user must define this value in the GUI during import. ***Important***: The alternative format assumes that the first timestamp corresponds with the first value in your fiber photometry data file.

**Figure 1.**
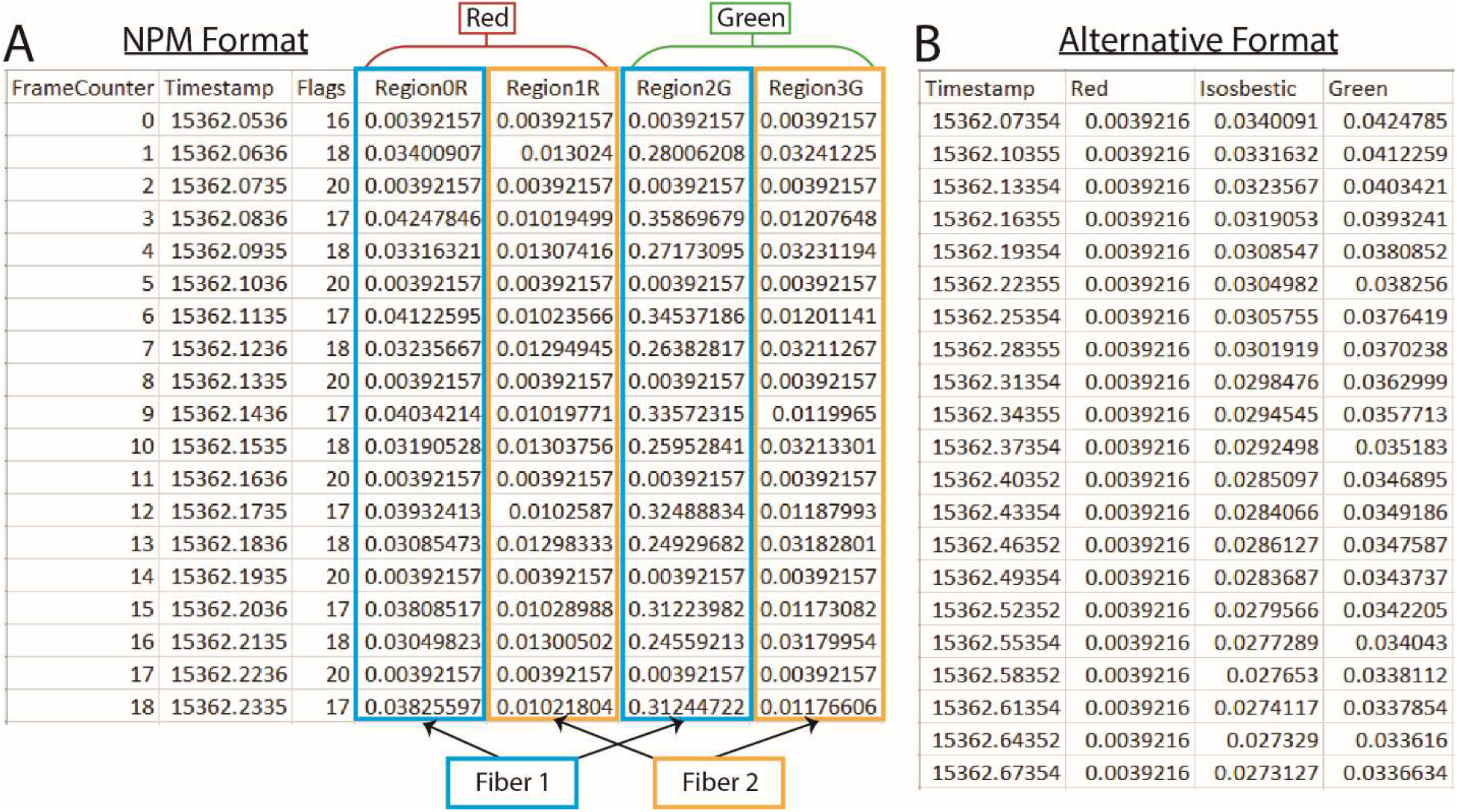
PhAT accepts two formats for photometry data. **A.** Example of an output csv file from Neurophotometrics (NPM). **B.** Example of alternate format photometry data csv file.

**Figure 2.**
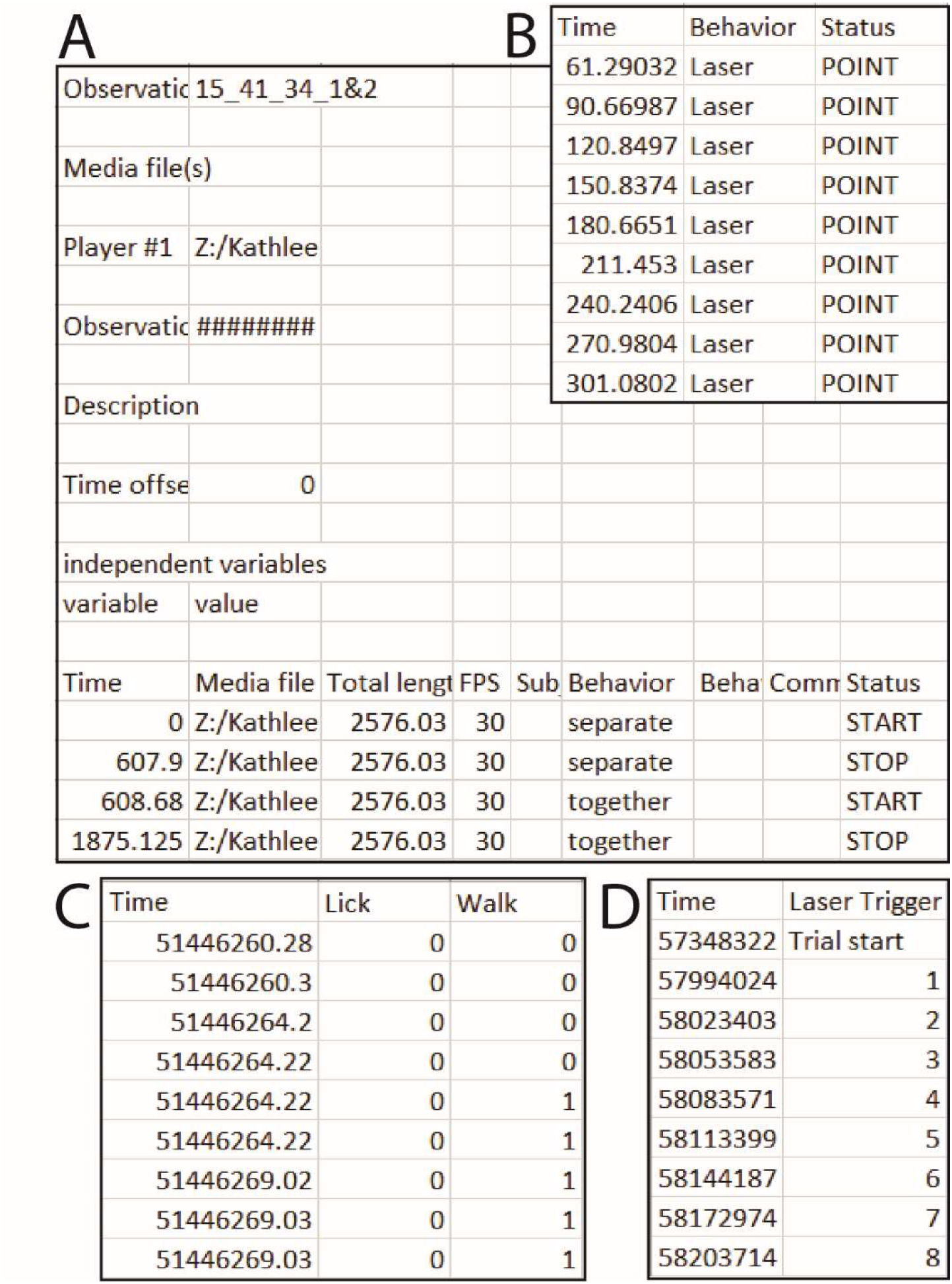
PhAT accepts two formats for event and/or behavior data. **A.** Example of an output csv file from BORIS. **B.** Example of a simpler format that will also work with the BORIS option in PhAT. **C.** One example csv file that will work with the Alternative format input option. In this example the user would enter “0” in the “value where behavior is not occurring” widget. **D.** A second example csv file that would work with the alternative format option. In this example “Trial start” would be entered for “value where the behavior is not occurring”.

### Protocol steps

1. Open the Graphical User Interface (GUI).
  a. Activate your virtual environment (see Alternative protocol 1, section 1d).

- Running with Jupyter Notebook (Option 1)
  b. If you would like to utilize Jupyter Notebook to deploy the server, simply navigate to the “FiberPho_Main” folder then run the “jupyter lab” command. Open the notebook (PhAT_gui_notebook.ipynb) file and begin to execute each cell (block of code) from the top, making sure to let each cell finish execution before continuing to the next.
  c. Upon execution of the last cell, a local URL will be displayed in the corresponding output cell that navigates to the GUI (e.g. http://localhost:####).

- Running with the Python Script (Option 2)
  b. In your terminal/command prompt, navigate to the location of the “FiberPho_Main” folder and run the following command (also listed at the top of the PhAT_gui_script.py file):
    “panel serve –show PhAT_gui_script.py –websocket-max-message-size=104876000 –autoreload”
  c. This command will launch the GUI in a new browser window or tab. To properly shutdown the GUI, press “Ctrl + C” on your keyboard.
  d. Any code changes made to the PhAT_gui_script.py file will refresh the entire server instance. To avoid this, omit the “--autoreload” argument.

2. Importing fiber photometry data. You will need to create an object for each recording from each fiber optic. Once these objects have been generated using the below steps, they can be re-imported for subsequent analysis via the “Reload Object” card on the left side of the GUI.
  a. Navigate to the “Create new fiber object” card at the top left corner of the GUI (Fig 3a).
  b. Click “Choose file” and select your fiber photometry data file.
  c. **(Option 1)** Working with Neurophotometrics (NPM) data
    i. Select the NPM output file (Fig 1a).
    ii. Enter the number of the fiber you wish to import from the file in the fiber number widget (see Materials for more in-depth explanation).
  c. **(Option 2)** Working with non-NPM data
    i. Select a .csv with photometry data in the alternative format (Fig 1b).
    ii. Uncheck the “Npm format” box.
  d. Enter the name of your fiber photometry object in the object name widget.
    i. Note: Use a long descriptive name without spaces as this name will be used as the main identifier for this data and will serve as the filename if the object is exported. We often use “Experiment_animalnumber_brainregion_sensor.”
  e. Optional but recommended: Enter descriptive information for your object, such as animal number, acquisition date, brain region, and sensor/fluorophores present, and experimental considerations (experiment disruptions, etc). These values will appear in the fiber data table to provide you with information on the experiment the data is associated with.
  f. Optional: Trim your data.
    i. Adjust the value in the “Exclude time from beginning of recording” box to specify how much time in seconds you would like to remove from the beginning of your file.
    ii. Adjust the value in the “Stop time from the beginning” box to specify the last value in seconds you would like to include in the trace. Leaving the value as −1 will not remove any time from the end of the file.
  g. Click the “Create Object” button.
    i. Your object has been created.
    ii. A successful creation will cause a green pop up in the bottom right corner of the GUI. The object’s information will be displayed in the table in the top right corner labeled “Display Object Attributes”.
3. Importing behavior data. This step is not required for all cards, but is necessary for any analysis that incorporates these data.
  a. Navigate to the “Import behavior” card at the top center of the GUI (Fig 3b).
  b. **(Option 1)** Using the BORIS format (Fig 2a/b)
    i. Make sure the BORIS format tab at the top of the card is selected.
    ii. Choose a fiber object from the drop-down menu.
    iii. Click “Choose file” and select your behavior data file.
    iv. Click “Import Behavior Data”.
    v. Your behavior is now saved with your fiber object.
  b. **(Option 2)** Using the Alternative format (Fig 2c/d)
    i. Select the Alternative format tab at the top of the card.
    ii. Choose a fiber object from the drop-down menu.
    iii. Click “Choose file” and select your behavior data file.
    iv. Select the time unit of your “Time” columns from the drop-down menu.
    v. Enter the value your file uses to signify when a behavior is not occurring. (This value would be “0” in the first example and “Trial Start” for the second (Fig 2c/d)).
    vi. Enter the minimum time you would like to use between bouts in the “time between bouts” box.
      1. This time should be in the same unit as the timestamps in your file.
      2. The start of each bout will have to be preceded by at least this amount of time in which the behavior is not occurring. For example if we use 0.5 secs for this value, any inter-bout interval < 0.5 sec will be considered part of the same bout but intervals > 0.5 sec will be considered two distinct bouts.
    vii. Click “Import Behavior Data”.
    viii. Your behavior is now saved with your fiber object.
4. Save fiber objects. Each fiber object you create in the GUI can be saved for later. This allows you to begin analysis, close the GUI, and reopen and import your objects without losing any progress.
  a. Navigate to ”Save fiber objects for later” card on the left hand side of the GUI (Fig 3a).
  b. Choose one or more fiber objects from the menu.
  c. Click the save object(s) button.
    i. The objects will be saved as a pickle (pkl) file. The filename will be the name of that object, and they will be saved into the “Fiberpho_Main” folder.
    ii. Once saved, the objects can stay in this folder or be moved to any other folder.
5. Reload fiber objects. If you’ve saved fiber objects as pickles using the “Save fiber objects” card, you can reimport them to resume an analysis using this card.
  a. Navigate to the “Reload saved Fiber Objects” card on the left-hand side of the GUI (Fig 3a).
  b. Click “choose files.”
  c. Navigate to and select all the .pkl files you would like to upload.
  d. Click the upload object(s) button.
    i. The software will confirm that the object was saved with the same version of the software you are using. If it is not, a warning pop up will appear in the bottom right corner of the GUI, and a message denoting the objects with potential incompatibilities will appear in the terminal. The object will still load but may cause errors when used with one or more cards.
    ii. A successful creation will cause a green pop up in the bottom right corner of the GUI. The object’s information will be displayed in the table in the top right corner labeled “Display Object Attributes”.
6. Combine fiber objects. You may want to combine two fiber objects either from the same file after cropping out a large artifact or to combine two files from the same trial or experiment. To do this you will create two fiber objects using the “Create new fiber objects” card and then combine them using the “Combine two existing fiber objects” card.
  a. Navigate to the “Combine two existing fiber objects card” on the left-hand side of the GUI (Fig 3a).
  b. Enter a name for your new object.
  c. Select the object you want in the beginning with the “First Object” widget.
  d. Select the object you want at the end with the “Second Object” Widget.
  e. Select how you would like to combine the times of each object using the “Stitch type” widget.
  f. Enter a time in the “x seconds” widget if you chose a stitch type that requires it.
  g. If you have successfully combined the fiber objects you should see a green box pop up in the bottom right hand corner after completion.
7. Delete fiber objects. Use the “Delete object” card to delete an object. This is particularly useful if you made a mistake importing/creating the object or adding behavior. No two objects can have the same name; trying to create a new object will not overwrite an existing object with the same name.
  a. Navigate to the “Delete object” card on the left hand side of the GUI (Fig 3a).
  b. Choose one or more fiber objects from the menu.
  c. Click the delete object(s) button.
8. Plot your data.
  a. Navigate to and expand the “Plot raw signal” card by clicking the green triangle on the left side of the card (Fig 3d).
  b. Choose one or more objects.
    i. An interactive graph will be made for each selected object. (See support protocol 2b for further instructions on interacting with graphs)
    ii. All traces associated with the object will be plotted together.
    iii. This tool can be useful for identifying large artifacts that you can then crop out (see step 2g) before recombining your data set (see step 6).
9. Normalizing your data. The “Normalize data” card will simultaneously linearize a trace by accounting for photobleaching and subtract motion artifacts to create the ΔF/F traces that are typically used in fiber photometry analysis. Because the most effective normalization strategy is often dependent on the experiment, we’ve created a flexible tool that allows you to normalize your data in different ways (Fig 4). Considerations for each option are detailed below.
  a. Navigate to and expand the “Normalize to a reference” card by clicking the green triangle on the left side of the card (Fig 3d).
  b. Choose one or more objects in the object selection box. Then click the “update options” button.
    i. Only channels present in each selected object will appear in the signal and reference dropdown boxes.
  c. Select the signal channel you wish to normalize.
  d. Select a signal-independent reference channel or “None” if you wish to skip the motion artifact removal step.
  e. Optional: Change the threshold for the goodness-of-fit for the biexponential fit
    i. Enter your desired threshold for an R^2 value. Fits that fall below the criteria will be ignored and your trace will be normalized to its median value instead of the biexponential decay
    ii. Set the threshold to 1 to skip the linearization-by-biexponential-fit step
  f. Choose a fit type for motion correction (Fig 4c/d).
    i. The difference between fit types is described in the considerations section below.
  g. Click the “Normalize Signal” button.
    i. This will normalize the signal and add the normalized signal to each object for future use.
    ii. The linear fitting process will be shown for each trace in a series of graphs to allow for a visual assessment of the fit. All the coefficients used to fit each channel will also be saved with the object.
    iii. If the goodness of fit for linearization is below threshold, the trace will be normalized to the median value of the trace, and you will be notified by a yellow warning pop-up and a message in the terminal.
    iv. The last of the graphs in the series will show the motion-corrected signal trace (via subtraction of the reference signal).

- Considerations for linearizing your trace Most of the time, you will want to linearize a trace by fitting to a biexponential curve (Fig 4a/b), which accounts for exponential photobleaching from the fluorophore as well as photobleaching of the patch cable, which may have different rates of decay. However, there are a few instances in which this is inadvisable, such as when you have no/little photobleaching, during very short recordings, or when your signal amplitude is greater than your photobleaching. The goodness of fit for your biexponential curve can be used to guide your decision of whether or not to linearize your trace via biexponential fitting.
- Considerations for subtracting motion artifacts The second step of the normalization process attempts to reduce motion artifacts by fitting your linearized signal trace to a linearized reference trace, such as the isosbestic channel or a channel corresponding to a non-sensor fluorophore (e.g. mCherry). As articulated in the *Strategic Planning* section above, the choice of ideal reference signal will depend on the specific sensor employed. You can skip this by setting the reference channel to none. The software provides two options for linearly fitting the reference to the signal for motion artifact correction both using the equation: Linfit = A_l_ Norm_ref_ + B_l_, with the differences stemming from how the coefficients are calculated. The first uses the “curve_fit” function from the python Scipy package to determine the coefficient A_l_ and B_l_ (Fig 4c), which relies on a non-linear least squares algorithm. The second option uses a linear fit algorithm we have coined a “quartile fit”. In this case A? = IQR_sig_/IQR_ref_ and B_l_ = Median_sig_ - A_l_ *Median_ref_. For the quartile fit, the reference is multiplied by the ratio of the signal interquartile range (IQR) to the reference IQR, so that the adjusted reference and the signal have the same IQR. Then that adjusted reference is shifted up or down so that its median is the same as the signal median (Fig 4d). Finally, we divide the linearized signal by the fitted reference to get the final normalized ΔF/F signal. Using the Least Squares option should be your starting point as it is the current standard in the field. However, there are instances in which this fails to eliminate clear motion artifacts, which are evident via simultaneous deviations in fluorescence in the signal and in the reference that are not eliminated by application of the “curve_fit” function (Fig 4c). In such instances, we recommend the quartile fit and subsequent visual inspection. Quartile fit is likely to be superior when you have large motion artifacts and/or small signals. We recommend using the same motion correction approach for all signal traces in the same experiment.

10. Visualizing behavior data.
  a. Navigate to and expand the “Plot Behavior” card by clicking the green triangle on the left side of the card (Fig 3d).
  b. Choose one or more fiber objects from the menu.
  c. Click update options.
    i. Only channels and behaviors found in all objects will appear in the menus.
  d. Choose any number of behaviors and channels.
  e. A graph will be created for each combination of object and channel with the selected behaviors overlaid as colored blocks (Fig 5).
11. Peri-event time series graphs. This card allows you to create a peri-event time series graph and save metrics from the analysis as results (Fig 6). This graph is the most common way to analyze fiber photometry data. It centers the signal around the beginning of particular events, such as all bouts of a particular behavior, so that signal changes can be averaged across multiple events. Our card allows you to graph either the % change in the signal or the Z-score with a user-defined baseline as appropriate for your experimental design.
  a. Navigate to and expand the “Peri-event time series plot” card by clicking the green triangle on the left side of the card (Fig 3d).
  b. Choose one or more objects in the object selection box, then click the “update options” button.
    i. Only channels and behaviors present in each selected object will appear in the signal and behavior widgets.
  c. Select the signal channel(s) you wish to visualize.
  d. Select the signal behavior(s) you wish to visualize.
    i. A unique graph will be created for each object, channel, and behavior combination.
  e. Enter the duration in seconds you would like plotted before and after the beginning of each behavior bout.
  f. Check the “Save CSV” box to save the dataframe used to make each plot as a csv.
  g. Check the “Use % of baseline instead of Zscore” box, to visualize the data as a percent change in the signal above your baseline instead of a z-score.
  h. *Optional: Choose a baseline for your z-score or percent change calculations. If you do not do this this, the baseline for each event will be the default option, “Each Event” (see below).
    i. Select the “baseline options” tab at the top of the card.
    ii. Select the region you would like to use as a baseline.
      - (DEFAULT) “Each Event” will use the entire time plotted for each bout, before and after the start of the behavior, as the baseline (Fig 6 blue regions/box).
      - “Start of Sample” allows you to select a time window at the beginning of your recording session to use as a baseline (Fig 6, purple region/box).
        i. Enter the time in seconds when your baseline period begins in the “Baseline Start Time” box.
        ii. Enter the time in seconds when your baseline period ends in the “Baseline End Time” box.
      - “Before Events” allows you to select a time window before each behavior bout to use as a baseline for that bout (Fig 6, dark pink).
        i. Enter the time in seconds when your baseline period begins relative to the onset of that behavior.
        ii. Enter the time in seconds when your baseline period ends relative to the onset of that behavior. (Ex. 8 seconds and 5 seconds will give you a three seconds baseline period that ends 5 seconds before the onset of each bout.)
      - “End of Sample” allows you to select a time window at the end of your recording session to use as a baseline for all of your behavior bouts (Fig 6, light pink).
        i. Enter the time in seconds *from the end of your recording* when your baseline period begins in the “Baseline Start Time” box.
        ii. Enter the time in seconds *from the end of your recording* when your baseline period ends in the “Baseline End Time” box. (Ex. 0 seconds will end your baseline period at the very end of your recording session.)
  i. *Optional: Reduce the number of events displayed on the graph, this will not affect the average or the csv if exported. This helps increase the speed in which graphs are created and make graphs with many events easier to interpret.
    i. Select the “Reduced displayed traces” tab at the top of the card.
    ii. Enter the first event you would like shown on your graph.
    iii. Enter the last event you would like shown on your graph. The default, -1, will choose the last event.
    iv. Enter the frequency of traces you would like displayed. (Ex. 3 will display every third trace)
  j. Click the “Create PETS plot” button.
    i. This simultaneously creates your peri-event time series graphs and a dataframe with some descriptive statistics for each plot that is stored in the corresponding object.
    ii. The Graph: Each trace will be plotted on the graph with the first events having the pinkest traces and the last events having the bluest traces. An average of all traces is plotted in black and the SEM is denoted by light gray shading. All traces can be toggled by clicking their name in the legend on the right. Double clicking the name will turn all traces beside the selected trace off.
    iii. The Results: Measures from each graph including the min and max amplitude, as well as the user input used to create the graph, will be stored in a results table within each object. These can then be exported using the “Export Results” card (see step 14)
12. Calculate Pearson’s R between traces. One benefit of fiber photometry and the Neurophotometrics system in particular is the ease with which simultaneous recordings can be collected in multiple channels, from multiple brain regions or across multiple animals. The time defined correlation card allows you to visualize and measure the Pearson’s correlation between two traces over a user-defined time window.
  a. Navigate to and expand the “Pearson’s Correlation Coefficient” card by clicking the green triangle on the left side of the card (Fig 3d).
  b. Choose one fiber object from each drop-down menu. They can be the same or different.
  c. Click “update options”.
  d. Choose a channel for each object from the widgets labeled “signal”.
  e. Declare the portion of your traces for correlation computation.
    i. Enter the start time in seconds in the “Start Time” widget. The default value of zero will use the beginning of the trace as the start of the window.
    ii. Enter the end time in seconds in the “End Time” widget. The default value of −1 will use the end of the trace as the start of the window.
  f. Click the “Calculate Pearson’s Correlation” button.
    i. Two graphs are created for each correlation, one that simply overlays each trace and a scatterplot showing the correlation and line of best fit.
    ii. The R value will be shown in the title of the graph, printed in the terminal, and saved in the corresponding results table stored with each object.
13. Calculate Pearson’s R during specific behaviors. The behavior correlation card works exactly like the time correlation card except that it compares all sections of each trace during which a specific behavior is occurring.
  a. Navigate to and expand the “Behavior Specific Pearson’s Correlation” card by clicking the green triangle on the left side of the card (Fig 3d).
  b. Choose one fiber object from each drop-down menu. They can be the same or different.
  c. Click “update options”.
  d. Choose a channel for each object from the widgets labeled “signal”.
  e. Select one or more behaviors from the behavior widget.
    i. A separate calculation will be performed for each behavior.
  f. Click the “Calculate Pearson’s Correlation” button.
    i. Two graphs are created for each correlation, one that simply overlays each trace and a scatterplot showing the correlation and line of best fit.
    ii. The R value will be shown in the title of the graph, printed in the terminal, and saved in the corresponding results table stored with each object.
14. Export results. The “Download Results” card allows you to export all the results from a specified analysis for multiple objects to a csv file (Fig 3d).
  a. Navigate to and expand the “Download Results” card by clicking the green triangle on the left side of the card.
  b. Enter a name for your results file in the “Output filename” widget.
    i. The type of analysis will be added to the end of the name for each file.
  c. Choose one or more objects from the “Fiber Objects” menu widget.
    i. Data for all objects will be combined into one file.
  d. Select one or more analyses from the “Result Types” menu.
    i. Each type of analysis will be exported into its own file.
  e. Click the “Download” button.
    i. Result csv files will be saved in the Fiberpho_main folder and can be moved anywhere once created.

**Figure 3.**
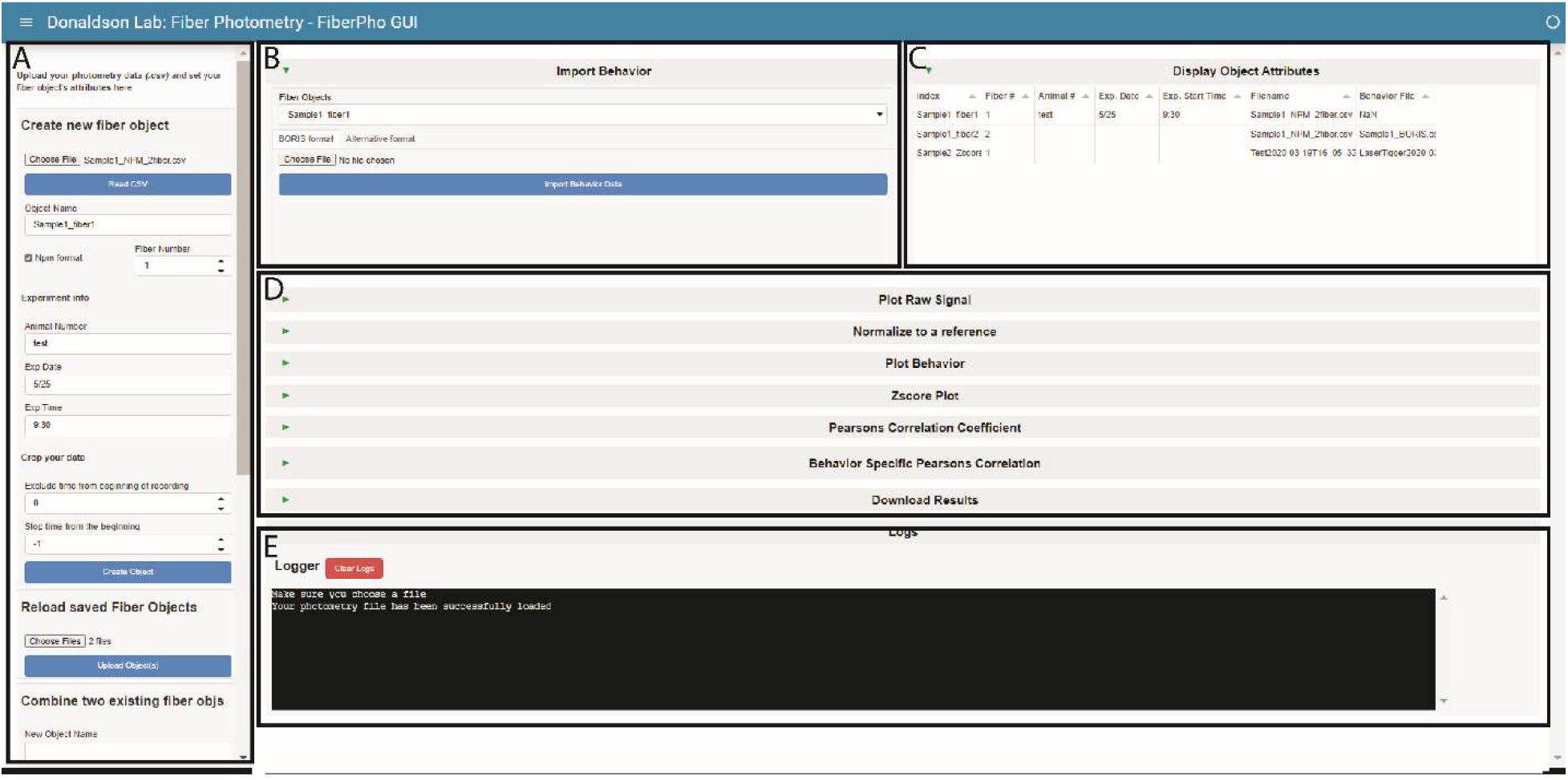
PhAT’s GUI layout. **A.** The sidebar. This houses all the cards that create, save or delete fiber objects. Use the respective scroll bar to access all the cards. **B.** The Import Behavior card. **C.** The Display Object Attributes table. This table will hold information on all the objects currently available in the GUI. **D.** This area holds all the cards available for analysis. They are all minimized in this figure, as denoted by the sideways green triangle. **E.** The Logger. This area is where information is shared with the user. It is also where all print statements will be output as well as in the terminal in the last output cell of the jupyter notebook.

**Figure 4.**
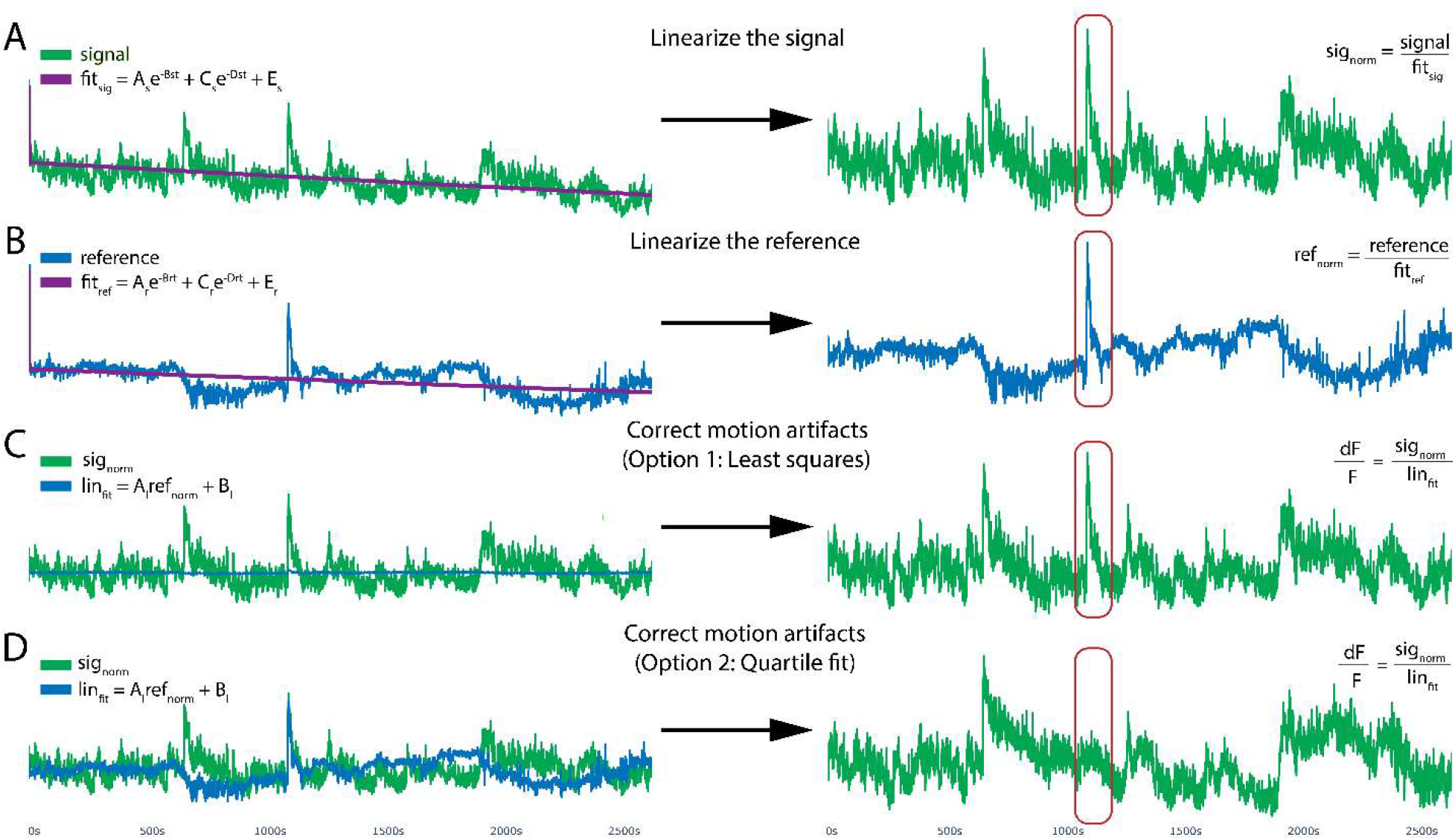
Motion reduction in PhAT. PhAT’s normalization card allows users to linearize their signal by removing the effects of photobleaching and reducing motion artifacts using one of two fitting algorithms. **A, B.** To optionally remove photobleaching, the program will fit a biexponential decay to your signal and reference traces and then divide by that fitted curve, resulting in the linearized signal (sig_norm_) shown on the right. **C.** The linearized reference (B) will then be fit to the linearized signal (A) to remove motion artifacts. This subfigure shows the reference (in blue) being fit to the signal using python’s built-in least squares algorithm. **D.** Reference fit using the alternative quartile fit algorithm, which in this specific case is more effective at removing the large motion artifact circled in red.

**Figure 5.**
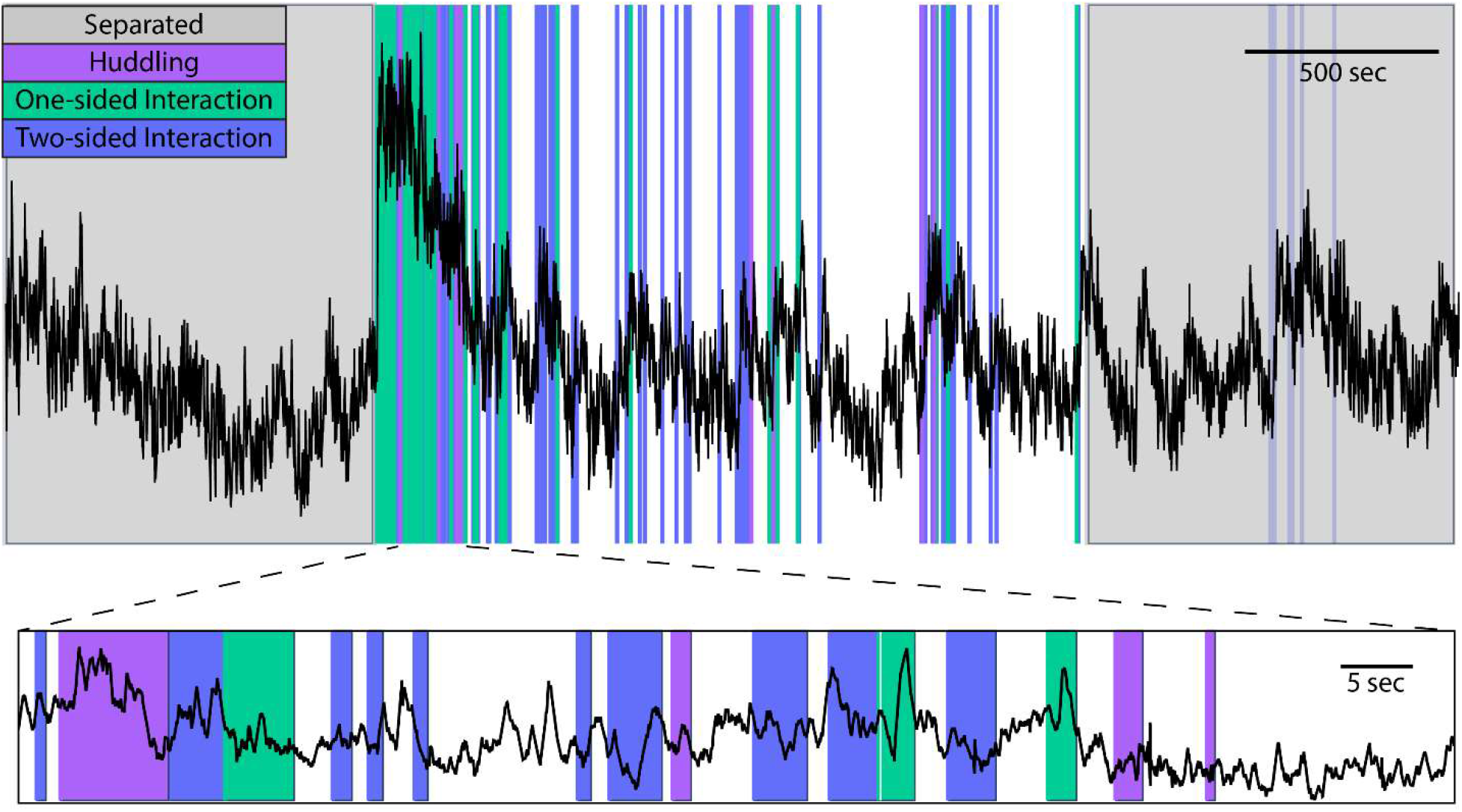
Example behavior plot generated by PhAT. PhAT’s plot behavior card allows you to visually represent any event data (colors) over your photometry traces (black). The interactive graphs allow the user to zoom in on regions of interest on the trace (a shown on bottom) to visually examine data and look for oddities and patterns before determining the best analysis strategies.

**Figure 6.**
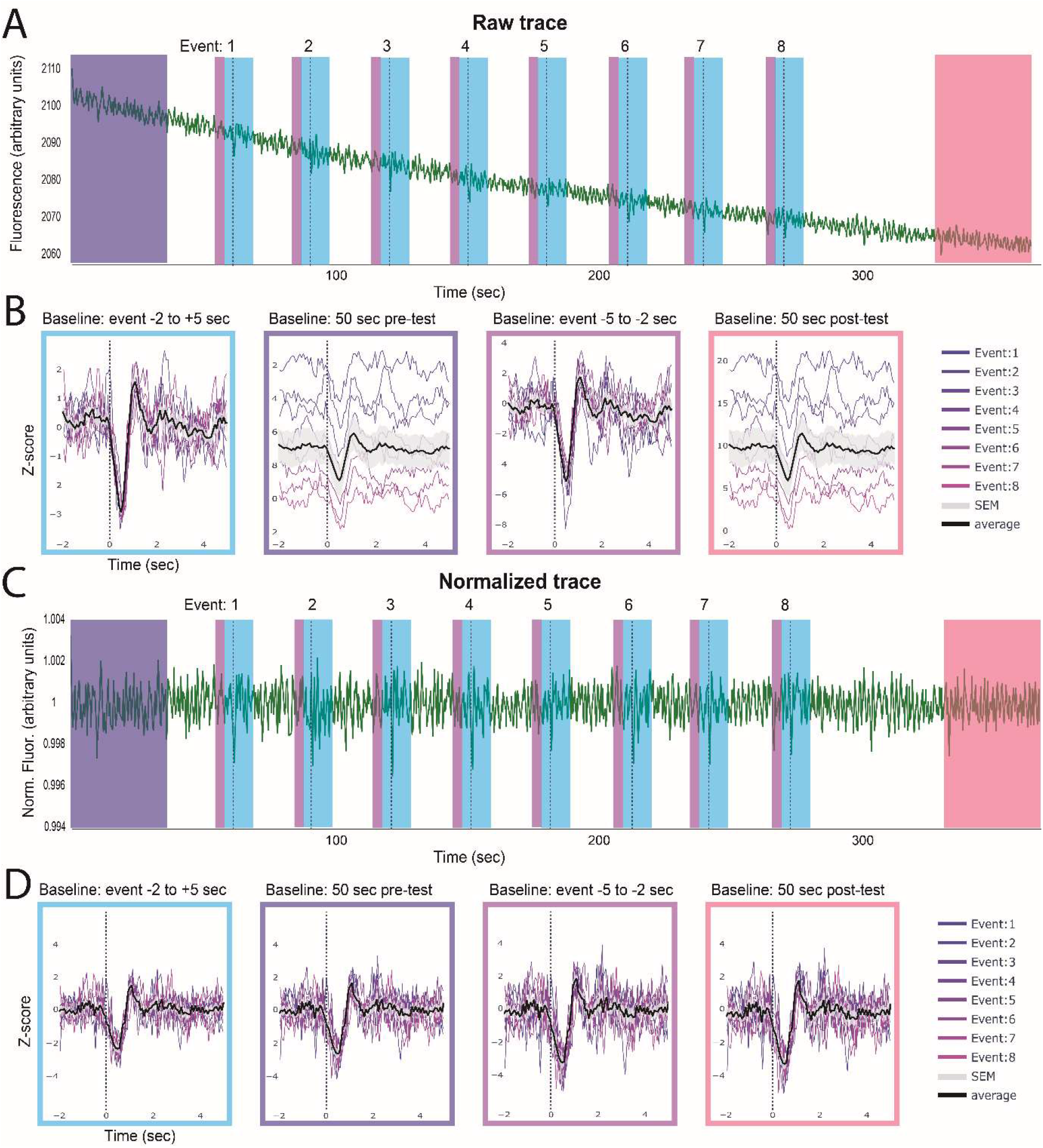
Identifying event-related changes in fluorescence. The peri-event time series (PETS) card allows the user to choose an ideal baseline for Z-scoring data. The above example shows GRAB_DA_-mediated fluorescence following optogenetic inhibition of the VTA (dotted line) **A.** The full trace with each individual event denoted by the dashed line. **B.** The peri-event time series plots with the z-scored trace using different baselines (indicated above each plot). The average fluorescence across events is shown in black with standard error in gray. **C.** The same data as in A but linearized and motion corrected. **D.** Peri-event time series on linearized trace using different baselines. **Summary.** These two examples show how choosing different baselines can affect the outcome of this analysis and the importance of linearization when using a baseline from the beginning or end of a session but not for event-adjacent baselines.

## Support Protocol 2a: Examining signal quality in your trace

Even after you have validated a sensor, there are factors that can cause poor signal in an animal or trial. This protocol is designed to help you evaluate signal quality for each experimental animal. Consider performing this analysis on an initial recording before deciding whether to include an animal in an experiment (Fig 7). This step does not require any behavior or event data, but it can provide additional valuable evaluation criteria.

**Figure 7.**
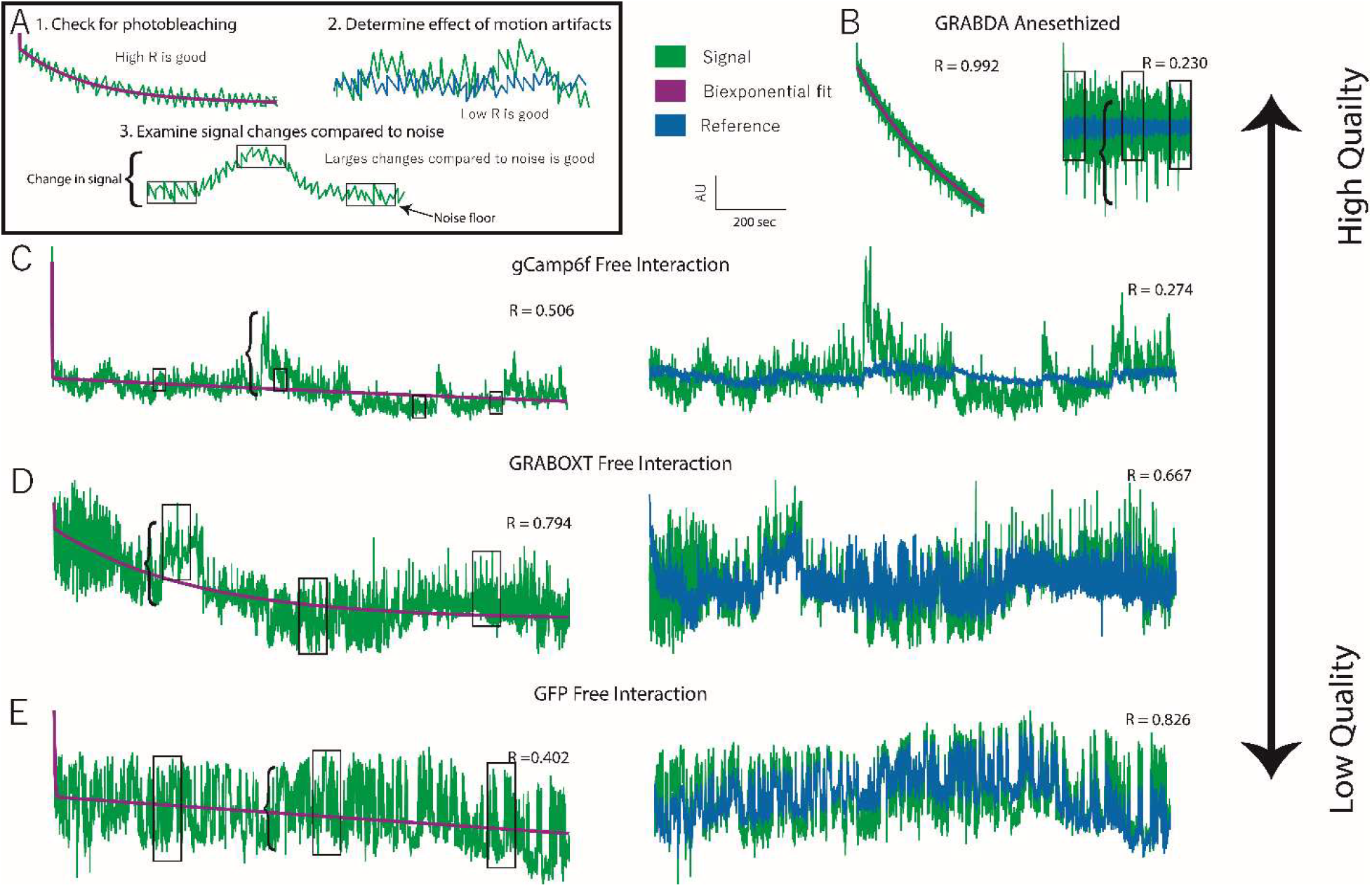
Data quality assessment. **A.** Examples of three features that indicate data quality shown with hypothetical data. 1. Evidence of photobleaching, which indicates presence of a fluorophore near that ferrule terminal. You can use the Pearson’s R value of a biexponential fit as an indicator of photobleaching. 2. Deviation in signal that is not present in the reference (i.e. a low signal:reference R value) indicates the presence of signal-based variation independent of variation due to motion artifacts. 3. Larger signal changes (bracket) relative to the noise floor (boxes) indicate good signal:noise ratio. **B.** Very high-quality data obtained from an anesthetized animal expressing GRAB_DA_ in the nucleus accumbens and receiving optogenetic inhibition of the ventral tegmental area. **C.** High quality data collected from a vole expressing GCaMP6f during social interaction. Evidence of photobleaching is moderate but other quality indicators are strong. **D.** Low quality data recorded from a vole expressing GRAB_OXT_ during social interaction. High signal:reference R-value indicates most variation is due to mation. **E.** Negative control data recorded from a vole expressing GFP in the prefrontal cortex during social interaction. Signal size:noise floor indicates low signal to noise and high r-vale for signal: reference indicate lack of signal independent of motion.

### Protocol steps

1. Assess the signal quality in your raw data.
  a. Plot your data using the plot raw data card (see protocol 2, step 6).
  b. Visually inspect the trace for evidence of photobleaching.
    i. Photobleaching should fit an exponential decay function, specifically a biexponential decay function (Fig 7a).
    ii. If your fluorophore is being expressed, there will likely be noticeable photobleaching when you begin recording from an animal.
  c. Look for variation in your signal.
    i. There should be small, uniform fluctuations in your signal which can be referred to as the noise floor (Fig 7a).
    ii. A good signal will also have large changes in the trace compared to the noise (Fig 7a).
  d. Do your fluorescent changes match what is known about the kinetics of your sensor?
    i. Every sensor has rise and decay constants, which determines how quickly changes in sensor fluorescence can occur.
    ii. Real changes in the signal can be slower than these constants, but faster changes must be caused by noise or motion artifacts.
2. Compare your signal and reference channels.
  a. Fit your signal to your reference channel using the normalization card (see Protocol 2, step 7).
    i. Evaluate similarities in channels by eye. If all large changes in your signal channel are also present in your reference channel, then those changes are due to motion and not due to changes in your sensor.
    ii. For a numerical measure of similarity, refer to the R-value printed in the title of the 5th panel, which indicates the correlation coefficient between the linearized reference and signal channels. A low correlation (<0.5) is a good indication that most changes in fluorescence are not due to motion artifacts. A high correlation (0.7 - 1) indicates significant motion artifacts but does not mean that there is not also a detectable signal because large motion artifacts may be overpowering differences between the channels. Effective motion correction can eliminate these artifacts and reveal a signal. If you choose to continue with these animals, it is important to repeat step 2.a.i with your normalized signal and critically evaluate any findings to confirm they are not due to motion artifacts (see next step).
3. Optional: Compare the peri-event time series graph of your normalized signal to your raw signal and reference traces using the peri-event time series card for any behavior that shows a reliable change in your normalized signal (Specific instructions in protocol 3).
  a. Confirm that this change is also detected and in the same direction in your raw signal. While the magnitude of the change and the noise may be different, but the general shape should replicate.
  b. Confirm that this change is not detected in your reference signal. Many behaviors are associated with a characteristic movement, which can cause consistent motion artifacts at the onset of your behavior. Because no normalization technique can eliminate all motion artifacts, it is important to be wary of any reliable signal change associated with a behavior *if you can also detect that signal change in the reference channel*.

## Support Protocol 2b: Interacting with graphs

All graphs in the GUI are created using the plotly module. The protocol below explains some basic ways to interact with the graphs (Fig 8). For more information you can view documentation at https://plotly.com/python/ or by clicking the navy square on the right end of the toolbar in the top right corner of each graph.

**Figure 8.**
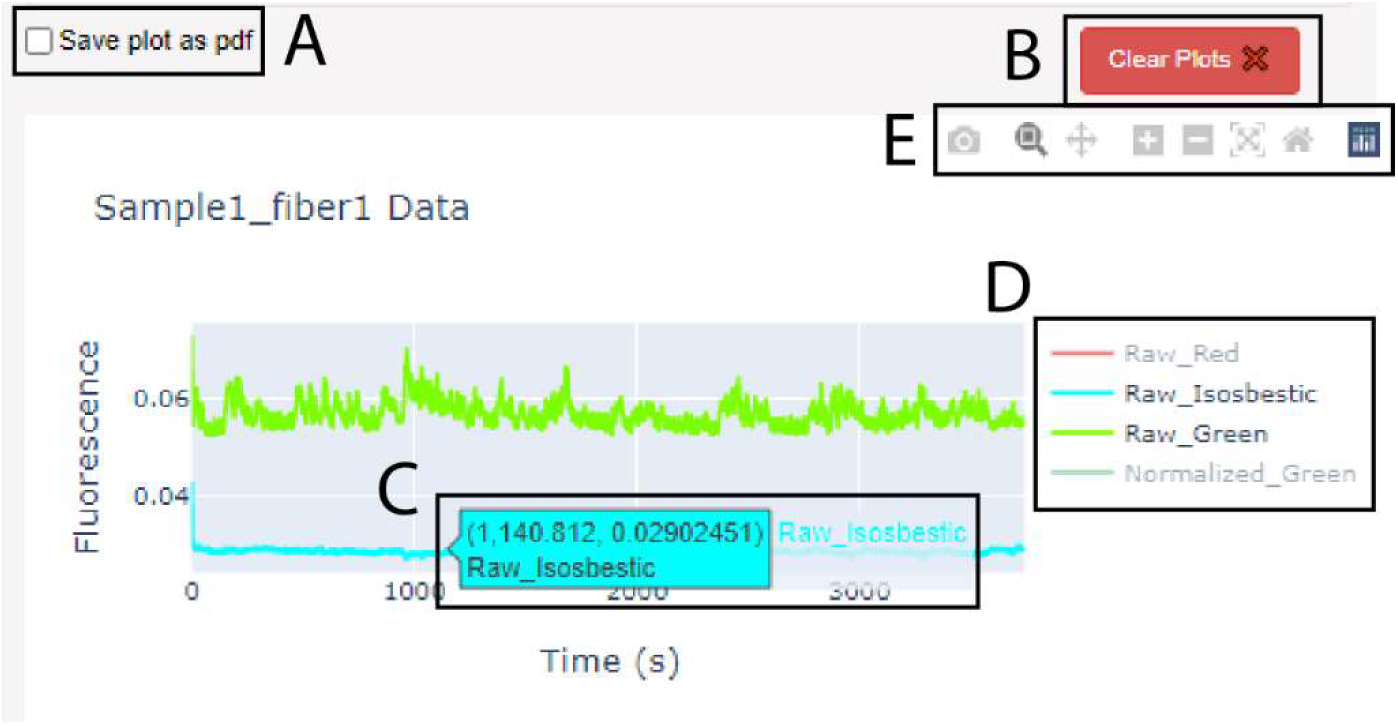
The graphs produced in PhAT are interactive. **A.** Checkbox widget used to save a graph as a pdf. Note: it must be checked before creating the graph. **B.** Clear plots widget used to delete the oldest/top graph in the corresponding card. **C.** Dialog box that appears when cursor hovers over a trace; values indicate x- and y-values for the trace at the cursor location. **D.** Graph Legend. **E.** Graph toolbar.

### Protocol steps

1. Save your graph as a pdf file.
  a. Check the “Save plot as pdf” checkbox in the bottom left corner of the corresponding card before creating the graph (Fig 8a).
  b. While this box is checked every graph you create will be saved in the fiberpho_main folder.
2. Delete a graph.
  a. Click the red “Clear plots” button in the bottom left corner of the corresponding card above the top plot (Fig 8b).
    i. Each click will delete the oldest (i.e. top) plot shown.
3. Identify a trace.
  a. Hover the cursor over a trace to open a dialog box with the raw data at the cursor location and the name of the trace as shown in the blue dialog box in Fig 8c. As needed, refer to the glossary for definitions.
    i. This is particularly useful for identifying timepoints for cropping traces.
4. Hide or display specific traces using the legend (Fig 8d).
  a. Click on a name in the legend to turn the name gray and hide that trace on the graph.
  b. Isolate a specific trace by double clicking the name in the legend, which will turn all other trace names gray and hide their traces.
5. Use the Toolbar (Fig 8e). (Note: The toolbar only appears when you hover over the graph with the mouse.)
  - The camera icon allows you to save the current view of your graph as a png to your downloads folder.
  - The magnifying glass allows you to zoom into a section of your graph by drawing a rectangle on the graph with your cursor.
  - The cross icon allows you to pan or move around the graph without changing the scale.
  - The plus and minus icons zoom in and out respectively with each click.
  - The X icon will auto scale your axes so the traces on the graph are maximized.
  - The home icon will reset your axes to the starting values.
  - The navy icon will direct you to the Plotly website where you can find other resources for creating and interacting with Plotly graphs.

## Basic Protocol 3: Adding Modules to PhAT

The modular and object-oriented structure of this software makes adding functionality straightforward for anyone familiar with python (Fig 9). This section outlines the overall design of the code and step-by-step instructions for adding new modules to the GUI. The alternative protocol explains how to work with fiber objects in the jupyter notebook, so that you can write and use new functions without adding them to the GUI.

**Fig 9.**
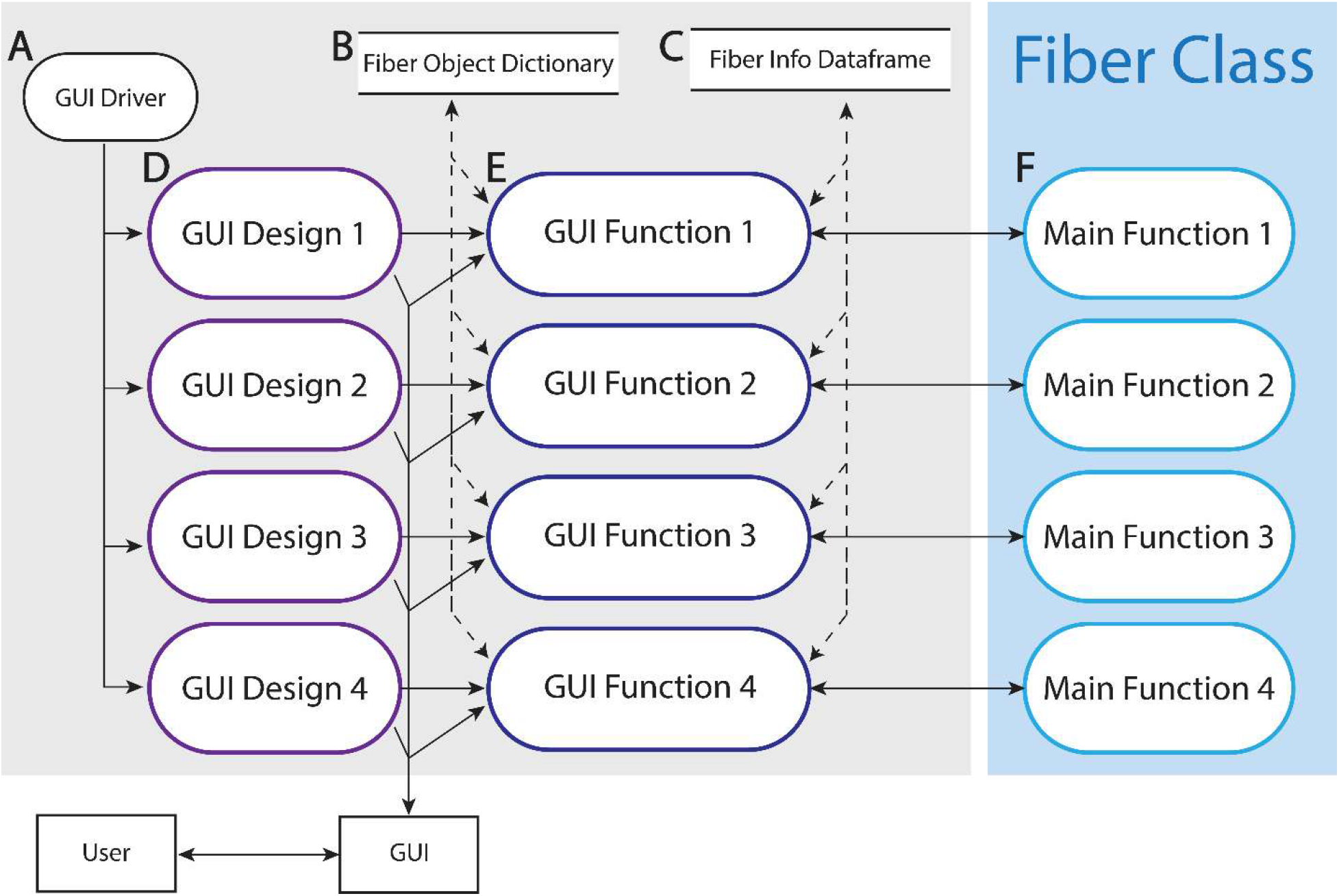
Data flow diagram of PhAT. The majority of the software is comprised of a series of modules denoted here as 1-4. Each module includes a section of GUI design, a GUI function and a main function. The gray box indicated components in the GUI files. The blue box indicates components in the FiberClass.py file. **A.** The GUI driver creates and displays the GUI. **B.** The fiber object dictionary is called fiber_obj and holds all available object using the object name as the key. **C.** The fiber info data frame holds some key attributes of each available fiber object to be displayed in the “Display Fiber Attributes” table. **D.** For an example of a GUI design section see “”#Plot raw signal Card” in PhAT_GUI_script.py. **E.** For an example of a GUI function see run_plot_traces in PhAT_GUI_script.py. **F.** For an example of a main function see plot_traces in FiberClass.py.

### Software Design

The code consists of 5 sections:

1. The fiber class (member function/or class function) This file holds the fiber class. It is where all the functions that work to visualize, manipulate and analyze your data are housed. These functions are all in the FiberClass.py file.
2. The import and initial declarations This section imports all the necessary packages and creates a dictionary that will hold all fiber objects using their obj_name as a key, and a data frame that holds basic information about each object to display to the user.
3. GUI definition In this section the cards seen in the GUI are designed. This includes declaring any user inputs that may be desired as well as adjusting the aesthetics of the GUI. This section is housed in the second half of the PhAT_gui_script.py or the second cell of the PhAT_gui_notebook.ipynb.
4. GUI functions These functions reformat the user input so that it can be used to call the respective fiber class functions. These functions also adjust the GUI to display outputs from the fiber class function. These functions can be found in the first half of the PhAT_gui_script.py or in the first cell of the PhAT_gui_notebook.ipynb.
5. GUI creation and serving This section adjusts any global attributes of the GUI’s design and then deploys the GUI for use.

#### Protocol steps

1. Create a function in the FiberClass.py file.
  a. Use this syntax for your function: def function_name(self, additional, arguments)
    i. Creating a function in a class is exactly like creating a regular function except that your first argument will always be the key word self, which will refer to the object you use to call the function.
    ii. Your arguments can be any user input as well as other objects if you would like to do an analysis that requires two or more objects.
    iii. In your function you will be able to access all attributes of the object you use to call the function and any objects you include as arguments
2. Access your Fiber objects.
  a. In the PhAT_gui_notebook.ipynb file or the PhAT_gui_script.py file: Objects will then be stored in the fiber_objs dictionary. The key for each object will be the obj_name. You can access your object using the code “fiber_objs[obj_name]”.
  b. In a fiberclass function: Use the code word “self”, to refer to the object you used to call the function. Any additional objects will just be referred to by their argument variable name.
3. Access fiber object attributes to use in your functions.
  a. Use dot notation to access variables stored within an object (attributes)
    i. The syntax is: object.attribute
    ii. All attributes of a fiber objects are described in table 2.
    iii. Examples:
      self.fpho_data_df
      fiber_objs[obj_name].channels
4. Create the GUI interface. If you would like to incorporate your new function into the GUI, you will need to make a new card for the GUI. All the cards use the panel holoviz package. For detailed documentation look here. https://panel.holoviz.org/reference/index.html#widgets
  a. Create an appropriate widget for each piece of user input you would like to collect.
    Some helpful widgets are:
      1. Fileinput
      2. Select_multiselect
      3. Textinput
  b. Create a button.
    i. When clicked the button will call a GUI function. Panel does not allow you to pass any arguments to said GUI function besides the number of times it was clicked (which is not typically valuable).
    ii. However, the GUI function will have access to all the user inputted values and any other variables defined in the file outside of other GUI functions. This includes the fiber_objs dictionary which holds all the objects and the fiber_data dataframe which holds some key attributes of each object.
  c. Organize all the widgets and buttons onto a card.
    i. You can align widgets in a row or column.
    ii. Then create a card with all your rows, columns and additional widgets or buttons.
    iii. For more detailed information. https://panel.holoviz.org/reference/index.html#layouts
  d. Optional: Use the existing gui layouts as a starting point.
    i. Example 1: “Create new fiber object”
    ii. Example 2: “Behavior Specific Correlation Plot”
5. Create the GUI function. The GUI function is used to connect the GUI to the fiberclass function.
  a. Access all the user input from the GUI.
    This can be done by accessing parameters of the widget. Most commonly you will simply use the syntax: widget_variable_name.value to get the value currently displayed in that widget. However, some widgets have multiple parameters that may be useful to access.
      1. for example. Fileinput() https://panel.holoviz.org/reference/widgets/FileInput.html
  b. Reorganize the user input so that it is compatible with your fiber class function.
    i. For example, if you allow the user to input multiple values for parallel processing using a widget like Multiselect, widget.value will return a list. You may want to iterate over that list.
    ii. Or you ask the user to pick a fiber obj by the obj_name variable, you will then have to actually access that object using the fiber_objs dictionary
  c. Call your fiber_class function.
  d. Update the GUI to display output from the function.
    i. The most common way I’ve done this is with plot_plane.
  e. Optional: Add try/catch phrases to ensure user input is valid before calling your main function.
  f. Optional: Use the existing gui functions as a starting point.
    i. Example 1: “def_upload_fiberobj”
    ii. Example 2: “run_plot_PETS”

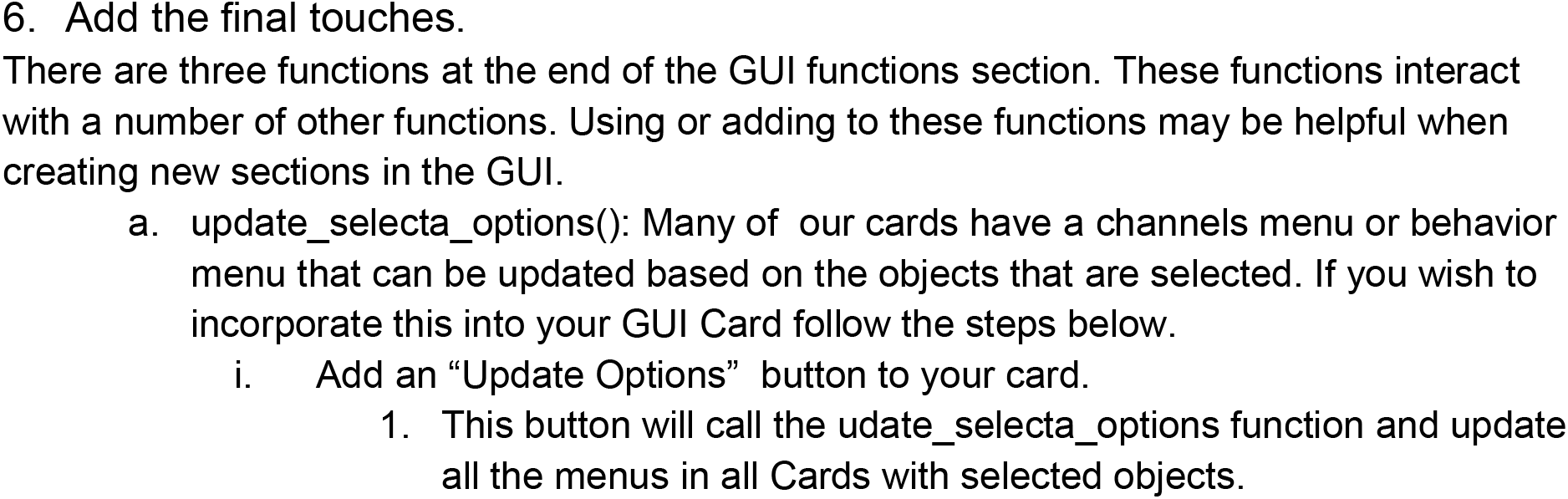
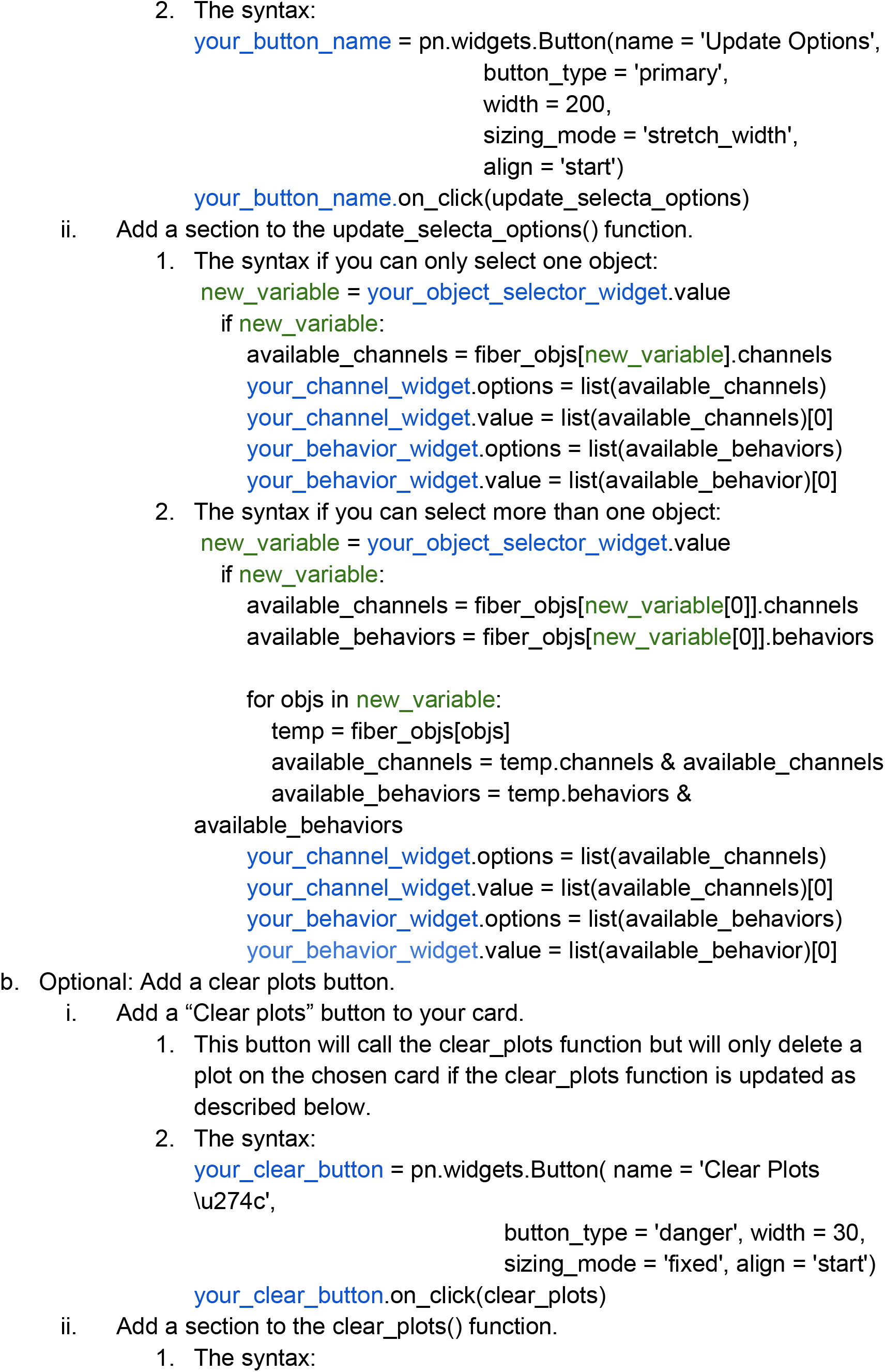
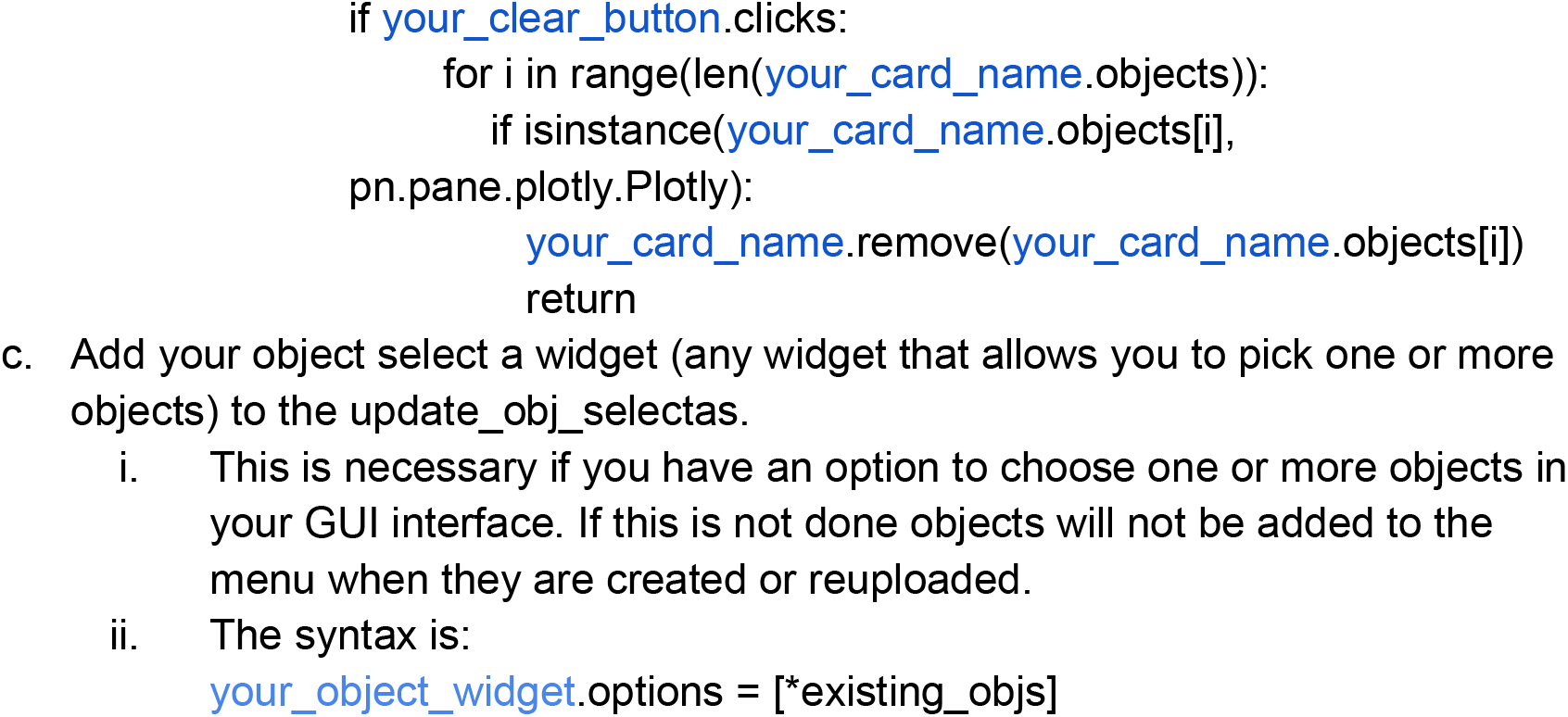

7. Update imports and the requirement.txt file with new packages.
  a. Add an import statement to the beginning of any file in which you are using a new modules/packages/libraries
  b. Optional: Add a line to the requirements.txt file in the PhAT folder for each new module, package, or library.
    i. This allows others using the code to easily install any new dependencies following protocol 1
    ii. Use the format: module name == version number

**Table 2:** Object Attributes. Here we list all the attributes of a fiber object, their data type a short description and the functions that modify them. All attributes are declared upon creation of the object and filled with an empty value if not provided.

| Attribute | Type | Description | Updated |
| --- | --- | --- | --- |
| obj_name | string | The name given to this object. Will be used to identify the object in the GUI and in any files exported from the GUI | N/A |
| fpho_data_df | Dataframe | A dataframe that holds all your photometry and behavior data. It has columns for time, each channel and each behavior. | normalize_a_signal<br>import_behavior_data |
| fiber_num | int | The fiber number this object corresponds to in the file. Only relevant for NPM file formats | N/A |
| animal_num | string | Optional* Defined by user input to give you additional information on the objects data | N/A |
| exp_date | string | Optional* Defined by user input to give you additional information on the objects data | N/A |
| exp_start_time | string | Optional* Defined by user input to give you additional information on the objects data | N/A |
| start_time | float | Time after the start of the photometry file that the object traces begin | N/A |
| stop_time | float | Time after the start of the photometry file that the object traces end | N/A |
| start_idx | int | Index of the start time in the photometry file | N/A |
| stop_idx | int | Index of the stop time in the photometry file | N/A |
| frame_rate | float | Frame rate of the photometry file | N/A |
| filename | string | The name of the csv file that your fiber photometry data was imported from | N/A |
| beh_filename | string | The name of the csv file that your behavior data was imported from | import_behavior_data |
| behaviors | set | All the behaviors that exist for this object | import_behavior_data |
| channels | set | All the channels that exist for this object | normalize_a_signal |
| sig_fit_coefficients | str | The coefficients A-E used to make the biexponential fit to the signal | normalize_a_signal |
| ref_fit_coefficients | str | The coefficients A-E used to make the biexponential fit to the reference | normalize_a_signal |
| sig_to_ref_coefficients | str | The coefficients A and B used to linearly fit the reference to the signal | normalize_a_signal |
| version | int | The version number of the object. This will only change if the software is updated to | N/A |
| color_dict | dict | A dictionary that determines the color associated with each channel for plotting | N/A |
| PETS_results | Dataframe | Houses a number of measures from your PETS analyses. | plot_PETS |
| beh_corr_results | Dataframe | Houses a number of measures from your behavior correlation analyses. | behavior_specific_pearsons |
| correlation_results | Dataframe | Houses a number of measures from your time correlation analyses. | pearsons_correlation |

## Alternative Protocol 3: Creating new functions for use in the jupyter notebook

Adding new functions to the GUI can make sharing those functions with other users easier and can decrease the time it takes to process your own data. However, it is not necessary to add a function to the GUI to run additional analyses on an object you’ve created and edited in the GUI. Below we describe how to create a new fiberclass function and how to access your fiber objects from the jupyter notebook and the attributes within that object.

### Protocol steps

1. Create a function in the FiberClass.py file.
  i. Use this syntax for your function: Def function_name(self, additional, arguments)
    i. Creating a function in a class is exactly like creating a regular function except that your first argument will always be the key word “self”, which will refer to the object you use to call the function.
    ii. Your arguments can be any user input as well as other objects if you would like to do an analysis that requires two or more objects. Ex. (beh_correlation)
2. Access your Fiber objects.
  a. In the PhAT_gui_notebook.ipynb file or the PhAT_gui_script.py file: Objects will then be stored in the fiber_objs dictionary. The key for each object will be the obj_name. You can access your object using the code “fiber_objs[obj_name]”.
  b. In a fiberclass function: Use the code word “self”, to refer to the object you used to call the function. Any additional objects will be referred to by their argument variable name.
3. Access Fiber object attributes.
  a. Use dot notation to access variables stored within an object (attributes)
    i. The syntax is: object.attribute
    ii. All attributes of a fiber objects are described in table 2.
    iii. Examples:
      self.fpho_data_df
      fiber_objs[obj_name].channels
4. Call your new function using the PhAT_gui_notebook.ipynb file. Now that the function is created in the fiberclass you can call that function directly from the PhAT_gui_notebookipynb.
  a. Call your fiberclass function using dot notation.
    i. You will still need to create your object(s) using the GUI.
    ii. The syntax for calling your obj will look like:
      fiber_objs[obj_name].my_new_function(all, of, my, arguments)
      Or my_obj = fiber_objs[obj_name] my_obj.my_new_function(all, of, my, arguments)

## COMMENTARY

### Background

Photometry approaches are rapidly becoming commonplace in systems neuroscience laboratories. Unfortunately, the technology that has enabled acquisition of fluorescent signals has outpaced toolkits for analysis of the resulting data. Many labs have developed in-house analytical solutions that cannibalize code from various groups; the result is a mish-mosh of approaches with limited opportunities for cross-platform/cross-lab validation or comparisons. PhAT provides a free, open-source GUI-driven platform that can integrate photometry data collected from systems generated by different manufacturers/labs. It requires no prior coding experience and enables bespoke data interrogation through the addition of new modules.

PhAT is not the only open-source fiber-photometry analysis software. GuPPY and pMAT both provide attractive alternatives (Sherathiya et al., 2021; Bruno et al., 2021). These packages also offer a handful of analyses that have yet to be included in PhAT, such as peak finding. In addition, pMAT uses Matlab for its operations, which for some labs may provide advantages based on local expertise. However, PhAT has a few major strengths that make it useful for a range of labs and applications. The software works directly with NPM data outputs and can also accept data from other sources. We provide multiple approaches for signal normalization, and straightforward and flexible visualization of traces to facilitate selection of an optimal normalization approach for a given dataset. Our flexible, object-oriented design makes module-addition straightforward. Of course, there are many potentially informative analyses that have not yet been incorporated into PhAT. We hope that community-driven module development will expand the utility and functionality of this software. Finally, PhAT includes options for cross-trace similarity comparisons, which are essential for quality assessment and enables novel interrogation of signals across brain regions or across animals. Thus, PhAT provides new features and a robust platform for expansion of photometry analyses.

In addition to flexible analytical solutions provided by PhAT, we have also provided information on experimental best practices for photometry. To our knowledge, no other resource succinctly addresses considerations related to sensor selection and validation, reference signals, experimental design, and optimal fluorescence detection. We also provide guidance on how to assess signal quality from individual recording locations/animals. Thus, this protocol extends beyond analytical software to improve the quality of data collected for photometry experiments, ideally improving scientific rigor and leading to more reproducible results.

### Troubleshooting and Critical Parameters

We have provided extensive information above related to experimental considerations that will ensure collection of relevant, high-quality data. We strongly encourage labs to take appropriate steps to ensure that they are acquiring high-quality, reliable fluorescence data prior to experiment initiation and extensive analysis.

As relates to PhAT, the most common issues arise from incompatibilities with supporting software. It is important to use the specified version of anaconda, jupyter notebook etc. Even so there can be times when there are still errors. In these cases, uninstalling and reinstalling the software or packages causing issues is a good place to start. You can also find assistance online. We have attached resources for troubleshooting these issues in the Internet Resources section below.

The most common errors in PhAT itself derive from incompatibilities with the software and the format of imported data. While checks exist to alert users of these errors, unexpected issues may occur. If there are issues using the GUI, first confirm that the data file you used contains data in the format described in the materials section of basic protocol 2. As with any software there will be bugs in the code itself. If you believe you have encountered an error in the software, please report it on the https://github.com/donaldsonlab/PhAT.

Finally, while no coding skills are required to use PhAT. If you decide to write your own code for module addition, then correct syntax is important.

### Statistical Analysis

PhAT calculates multiple metrics from event-related z-scores and percent change in ΔF/F, including the maximum and minimum values, the times at which these occur relative to the event, and the average change after an event. In addition, you can calculate the Pearson correlation coefficient for any two traces, which can be used to assess sensor signal quality, examine relationships across brain regions, and/or across brains. These metrics are calculated for each object or object pair individually, and subsequent group-level analyses should be carried out on the exported values using your preferred statistical analysis software.

### Time Considerations

Basic protocol 1: 20 minutes to 1 hour.

Alternative protocol 1: Less than 30 minutes.

Basic protocol 2: Varies depending on the amount of data and number of analyses you wish to do. Estimated 0 minutes to 3 hours.

Support protocol 2a: 10 minutes.

Support protocol 2b: 30 minutes to 1 hour depending on the quality of your data.

Basic protocol 3: 30 minutes or more depending on your familiarity with python and the complexity of the analyses you wish to add.

## DATA AVAILABILITY STATEMENT

The data that support the protocol are openly available in the Donaldson Lab Github repository at http://doi.org/10.5281/zenodo.7644327, **in folder** “sample data”.

## ACKNOWLEDGEMENTS

This work was supported by funds from the Dana Foundation, Whitehall Foundation, and NIH DP2MH119427 to ZRD and F30MH126607 to KZM. We would like to thank the members of the Donaldson Lab for provided data used to test PhAT, along with critical feedback on the manuscript. Mostafa El-Kalliny contributed coding advice and code review. Kelly (Emory University) and Kozorovitskiy (Northwestern University) Labs, and Emma Tinkham contributed to beta-testing. Yulong Li provided plasmids for GRAB_DA_ and GRAB_OXT_. Sage Aronson (Neurophotometrics) provided advice and insight into data quality metrics. We also thank the animals used to generate this data and the animal care staff at the University of Colorado Boulder.

## INTERNET RESOURCES

To access our code base visit: https://github.com/donaldsonlab/PhAT

For information on how to **install python** and relevant download links visit: https://www.python.org/downloads/ or https://wiki.python.org/moin/BeginnersGuide/Download

For information on how to **install anaconda** and the relevant download links visit: https://docs.anaconda.com/anaconda/install/

For information on how to **install pip** and the relevant download links visit: https://pip.pypa.io/en/stable/installation/

For information and tutorials on how to use **jupyterlab** or **jupyter notebook** visit: https://www.tutorialspoint.com/jupyter/jupyterlab_overview.htm or https://www.tutorialspoint.com/jupyter/jupyter_notebook_introduction.htm

For information on **Panel** the library used to construct the GUI visit: https://panel.holoviz.org/index.html

For information on **Panel’s Cards** and other layouts specifically, visit: https://panel.holoviz.org/reference/index.html#layouts

For information on **Panel’s widgets** specifically, visit: https://panel.holoviz.org/reference/index.html#widgets

For information on the way **Panel’s graphs** specifically, visit: https://panel.holoviz.org/reference/panes/Plotly.html

For general information on how to create and interact with **Plotly** visit: https://plotly.com/python/

